# *LPA* and *APOE* are associated with statin selection in the UK Biobank

**DOI:** 10.1101/2020.08.28.272765

**Authors:** Adam Lavertu, Gregory McInnes, Yosuke Tanigawa, Russ B Altman, Manuel A. Rivas

**Affiliations:** Biomedical Informatics Training Program, Stanford University, Stanford, CA 94305, USA; Departments of Bioengineering, Genetics, and Medicine, Stanford, CA 94305, USA; Department of Biomedical Data Science, Stanford University, Stanford, CA 94305, USA

## Abstract

Genetics plays a key role in drug response, affecting efficacy and toxicity. Pharmacogenomics aims to understand how genetic variation influences drug response and develop clinical guidelines to aid clinicians in personalized treatment decisions informed by genetics. Although pharmacogenomics has not been broadly adopted into clinical practice, genetics influences treatment decisions regardless. Physicians adjust patient care based on observed response to medication, which may occur as a result of genetic variants harbored by the patient. Here we seek to understand the genetics of drug selection in statin therapy, a class of drugs widely used for high cholesterol treatment. Genetics are known to play an important role in statin efficacy and toxicity, leading to significant changes in patient outcome. We performed genome-wide association studies (GWAS) on statin selection among 59,198 participants in the UK Biobank and found that variants known to influence statin efficacy are significantly associated with statin selection. Specifically, we find that carriers of variants in *APOE* and *LPA* that are known to decrease efficacy of treatment are more likely to be on atorvastatin, a stronger statin. Additionally, carriers of the *APOE* and *LPA* variants are more likely to be on a higher intensity dose (a dose that reduces low-density lipoprotein cholesterol by greater than 40%) of atorvastatin than non-carriers (*APOE*: *p(high intensity)* = 0.16, OR = 1.7, *P* = 1.64 × 10^−4^, *LPA*: *p(high intensity)* = 0.17, OR = 1.4, *P* = 1.14 × 10^−2^). These findings represent the largest genetic association study of statin selection and statin dose association to date and provide evidence for the role of *LPA* and *APOE* in statin response, furthering the possibility of personalized statin therapy.

## Introduction

The field of pharmacogenetics focuses on the intersection between pharmacology and genetics, with the goal of tailoring an individual patient’s drug regimen to both environmental and genetic conditions^1^. However, the majority of modern prescribing practices do not have the information necessary to implement or simply disregard pharmacogenetic recommendations, instead prescribing medicine is often a trial-and-error approach^2,3^. Once a patient is diagnosed with a particular disease, the physician typically prescribes the first-line therapy for that illness. That decision can be influenced by the severity of the illness, as in hypercholesterolemia (also known as high cholesterol) where an individual patient’s cholesterol levels help determine whether a lower or higher power statin is prescribed^4^. Additionally, patients may not respond to a particular drug and/or experience severe side effects leading them to return to the physician for adjustment or substitution. In this way, disease severity, drug efficacy, and side-effects are three factors that contribute to the creation of sub-cohorts of patients diagnosed with the same illness but who are on a second-line drug. Pharmacogenetics has not yet been widely adopted into clinical practice, but nonetheless influences patient care. Unbeknownst to the treating clinician, genetics can lead to variability in response to low efficacy or side effects, which may result in a change in patient care. Through the use of large cohorts of patients with both prescription and genetic data available, we may be able to identify a relationship between drug choice and genetics.

Statins represent an excellent opportunity for testing our ability to identify the relationship between drug choice and genetics for several reasons, (1) Statins are widely used to reduce low-density lipoprotein (LDL-c) levels in patients with increased risk of cardiovascular disease^4^. The prevalence of cardiovascular disease and hypercholesterolemia means that a study focused on statins should be well powered to detect relevant genetic differences within a population receiving statin therapy. (2) Prior research has already identified pharmacogenetic relationships for individual’s response to statin therapy, with regards to both efficacy and adverse drug reactions^5^. (3) Current guidelines from the National Institute for Health Care Excellence (NICE) in the UK specifically recommend that physicians titrate statin therapy to the maximum licensed or tolerated dose^4^. Additionally, there are several types of statins with a range of intensity which have varying degrees of efficacy in certain patients. Assuming physicians in the UK follow these guidelines, the result should be a population of individuals receiving statin therapy that have been titrated to their maximum dose. The selected statin and maximum tolerated statin dose represents the pharmacogenetic phenotype of interest to this study.

The UK Biobank offers an unprecedented opportunity to discover relationships between drugs and genetics^6,7^. Rich phenotype and genotype data is available for nearly 500,000 subjects, including prescription drug status at the time of subject enrollment. Phenotype data consists of ICD-10 coded diagnoses from the National Health Service (NHS), longitudinal clinical records from the NHS, and self-reported phenotypes collected during an enrollment survey. The UK Biobank offers two sources of drug data: longitudinal prescription records from the NHS for 230,000 participants as well as what medications participants were taking on the day they did their enrollment survey^8^. The former provides longitudinal data for drug usage over time, while the latter provides a snapshot of medication usage on a single day.

Previous studies of pharmacogenetics in the UK Biobank have determined allele frequencies in important pharmacogenes^9^ and associations between drug dose and drug side effects with pharmacogenetic phenotypes^10^. Statins were included in these studies, but these studies were focused on a narrow set of pharmacogenes, not the entire genome. There are 62 genes that have been reported to influence statin response with some level of evidence in PharmGKB^11^. PharmGKB levels of evidence are manually assigned by expert curators and consist of 4 levels, with the top two levels split into A (higher) and B (lower) subgroups. Level 1 annotations indicate a high level of evidence for the variant-drug combination, while Level 2 is moderate evidence, Level 3 is low evidence, and Level 4 is preliminary evidence^11,12^. Among these is *SLCO1B1*. Individuals carrying poor functioning alleles of *SLCO1B1* are at higher risk for simvastatin-induced myopathy^13^, which lead the Clinical Pharmacogenomics Implementation Consortium (CPIC) to create a dosing guideline for simvastatin prescribing based on *SLCO1B1* genotype^14^. Statin response in previous UK Biobank studies has only been studied in the context of *SLCO1B1* pharmacogenetics^9,10^.

We utilized the medication data collected during the UK Biobank subject enrollment process to identify cohorts of individuals receiving different types of statin therapy. We ran several GWAS on this cohort by using individuals on a particular statin as cases and the remaining individuals receiving other forms of statin therapy as shared controls, a method we call a drug selection genome-wide association study (DS-GWAS). We further interrogated significantly associated variants by assessing the association between carrying an alternate allele and statin dosage.

## Methods

### Population stratification in UK Biobank Data

We downloaded the non-imputed genotype data from the UK Biobank^6,7^. To reduce the impact of potential confounders due to population structure, we filtered the genotyping data cohort based on sample descriptions provided by the UK Biobank (Resource 531). Specifically, we removed samples that did not meet any of the following criteria: samples were of white British ancestry (“in_white_British_ancestry_subset”), used for the UK Biobank PCA calculation (“used_in_pca_calculation”), not an outlier in rates of heterozygosity or missingness (“het_missing_outliers”), did not have more than ten putative relatives (“excess_relatives”), and were not putatively sex chromosome aneuploids (“putative_sex_chromosome_aneuploidy”). This resulted in 337,138 unrelated individuals analyzed in this study.

### Variant annotation and quality control

We annotated the directly-genotyped variants using the VEP LOFTEE plugin (https://github.com/konradjk/loftee) and variant quality control by comparing allele frequencies in the UK Biobank and gnomAD (gnomad.exomes.r2.0.1.sites.vcf.gz) as previously described^15,16^. We used dbSNP (v137) to annotate rsIDs. For directly-genotyped variants, we focused on variants outside of the major histocompatibility complex (MHC) region (hg19 chr6:25477797-36448354) and applied the following filtering criteria^16^:

- The missingness of the variant is less than 1%, considering that two genotyping arrays (the UK BiLEVE array and the UK Biobank array) cover a slightly different set of variants^7^.
- Minor-allele frequency is greater than 0.01%, given the recent reports casting questions on the reliability of ultra low-frequency variants^17,18^.
- Hardy-Weinberg disequilibrium test p-value is less than 1.0× 10^−7^
- Manual cluster plot inspection. We investigated the cluster plots for a subset of variants and removed 11 variants that have unreliable genotype calls^16^.
- Passed the comparison of minor allele frequency with the gnomAD dataset as described before^16^.

### Statin selection genome-wide association study

We applied logistic regression with the firth-fallback option using a generalized linear regression model implemented in PLINK v2.00aLM^19^. With firth-fallback option, we apply the logistic regression by default and switch to bias-reduced Firth regression using a port of logistf() (https://cran.r-project.org/web/packages/logistf/index.html) whenever the logistic regression failed to converge or one of the cells in 2 × 2 allele count by case/control contingency table is zero. We included age (integer, computed based on the year of birth field in UK Biobank [UK Biobank Field ID: 34]), sex (0=female, 1=male), array (0=UK BiLEVE Axiom Array, 1=UK Biobank Axiom Array), and the first 10 genomic principal components of each individual were included as covariates in the GWAS. We used the Bonferroni corrected genome-wide significance threshold of 6.3× 10^−8^, based on the number of variants included in the analysis. Associations with a standard error of log-odds ratio greater than 0.2 were excluded from variant analysis. All positions and reference alleles were canonicalized with respect to the NCBI GRCh37 reference and the alternate alleles were treated as effect alleles.

We created a new GWAS framework for this study, a drug selection genome-wide association study (DS-GWAS). We used DS-GWAS to test for genetic associations related to selection of a particular drug within a single class of drugs. The DS-GWAS follows the traditional case-control GWAS paradigm, but tests subjects taking an individual drug in a drug class against subjects taking other drugs within the drug class (Fig. 1A). The study design follows a shared control design, where subjects will serve as both cases and controls depending on which drug is being tested (Fig. 1B)^20^.

**Figure 1.**
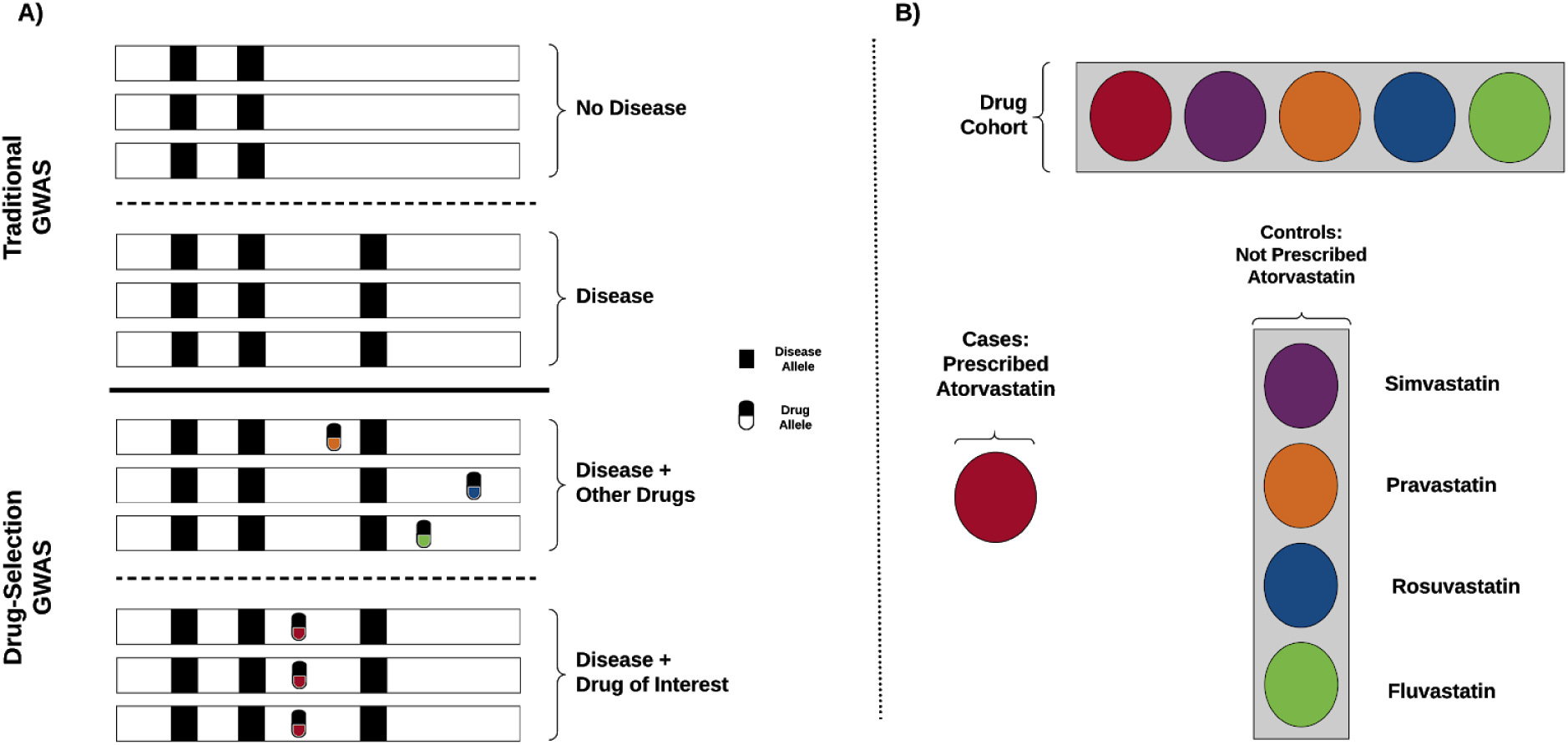
Drug-Selection GWAS (DS-GWAS) design and shared control framework. (A, top) Diagram layout of a traditional GWAS that aims to discover disease alleles through association with the disease phenotype of interest. (A, bottom) DS-GWAS design, controls for disease alleles with the goal of elucidating alleles associated with a specific drug. (B) Shared control study design for statins, the shared controls are composed of all individuals not on a particular drug (gray box), while cases are defined as individuals on a particular drug (circles). This figure demonstrates the DS-GWAS setup for atorvastatin.

We used DS-GWAS to test the five most frequently used statin drugs in the UK Biobank for drug-specific genetic associations using subjects taking each individual drug as the case cohort and all other subjects taking any other statin as the control cohort. The statins included were atorvastatin, fluvastatin, pravastatin, rosuvastatin, and simvastatin.

We constructed the statin cohort based on the UK Biobank individual level medication codes from the NHS, Data-Field 20003. Drugs included in the HMG CoA reductase inhibitors class (C10AA) of the Anatomical Therapeutic Chemical (ATC) classification, were used as inclusion criteria for the statin case cohort. The final list of included statins and their matched codes from UK Biobank Data-Coding 4 can be found in supplementary table 1. The case/control cohort definitions can be found in the quality control section of the results (Table 1). These cohorts are based on the self-reported medication data, which provides a snapshot of medication use at the time of enrollment.

**Table 1.** Case-control definitions and counts for GWAS cohorts used in this study.

| Case | Control | # Cases | # Controls |
| --- | --- | --- | --- |
| Hypercholesterolemia subjects | All subjects without hypercholesterolemia | 59,620 | 277,518 |
| All statin users | All subjects not on any statin | 58,112 | 279,026 |
| Simvastatin users | All subjects on a statin other than simvastatin | 30,923 | 13,851 |
| Atorvastatin users | All subjects on a statin other than atorvastatin | 11,302 | 33,472 |
| Rosuvastatin users | All subjects on a statin other than rosuvastatin | 1,959 | 42,815 |
| Pravastatin users | All subjects on a statin other than pravastatin | 1,426 | 43,348 |
| Fluvastatin users | All subjects on a statin other than fluvastatin | 124 | 44,650 |

We evaluated the association between statin selection genetics and the genetics of hypercholesterolemia. To do this we calculated the Pearson correlation coefficients between Wald Z-scores for all significant variants from a hypercholesterolemia GWAS with the corresponding values from a statin GWAS with all non-statin users as controls, as well as the correlation between z-scores from the hypercholesterolemia GWAS with the simvastatin DS-GWAS and atorvastatin DS-GWAS. The hypercholesterolemia cohort included all individuals who self-reported as having hypercholesterolemia (1473 - UK Biobank Data-Coding (UKBDC) 6) and/or were NHS coded as having hypercholesterolemia (E780 - UKBDC 19)^21^. We included all variants that were significantly associated with hypercholesterolemia in the analysis (*P* = 6.3 × 10^−8^).

Top hits from the DS-GWAS were evaluated for other pleiotropic associations using the PheWAS scan displayed in Global Biobank Engine variant page^22^.

### Genotype association with drug dose

We investigated the relationship between allele dosage of top hits from DS-GWAS and drug dosage. We used the primary care data to extract longitudinal prescription data for each query drug^8^. For all subjects on the drug, we calculated the prescribed daily dose by determining the average milligrams of drug per day for the last five prescriptions in the record. Individual prescriptions with a dose quantity two standard deviations away from the mean quantity were excluded. We required subjects to receive a minimum of five prescriptions of a drug. We then identified patients on a differing intensity levels (e.g. moderate vs high intensity dose, where high intensity is defined as a treatment that reduces LDL-c reduction greater than 40%) for each drug using the NICE guidelines^4^. This included simvastatin (low and moderate vs high (>=80mg)), atorvastatin (moderate vs high (>=20mg)), rosuvastatin (moderate vs high (>=10mg)), and fluvastatin (low vs moderate (>=80mg)). Pravastatin was excluded because all approved doses are considered low intensity^4^. Due to the small number individuals who carried homozygous alternate alleles for each variant, we binned all alternate allele carriers together. Then, we calculated the proportion of individuals on a high intensity statin and tested for a difference between homozygous reference individuals and alternate allele carriers using a test of equal proportions. Specifically, we computed a two-sided P value for a two-proportion z-test pooled for H0: p1 = p2 where p1 is proportion of non-carriers on a high intensity statin and p2 is the proportion of alternate allele carriers on a high intensity statin using the prop.test function in R with default parameters^23–25^.

### Known statin pharmacogenetic analysis

We evaluated the significance of pharmacogenetic variants known to be associated with statin response. The variants used were extracted from PharmGKB annotations file (https://www.pharmgkb.org/downloads, annotations.zip). Any individual variant who’s metadata mentioned an association with statin or HMG-CoA reductase was included in the analysis. Star alleles were excluded. For each variant we looked for associations with statin selection and statin dose. We determined the minimum p-value across each statin DS-GWAS to identify any statin selection associations with each variant, and determined the association between genotype and being on a high intensity regimen, as described in the previous section.

## Results

### Statin selection genome-wide association study

We included 337,138 participants who’s genotype data passed quality control metrics. Individual GWAS were run for users of each statin of five statins against all other statin users in the UK Biobank following the DS-GWAS framework. Additional GWAS were applied for use of any of the five statin and hypercholesterolemia. Final cohort sizes for each of the statin drugs can be seen in Table 1.

We identified seven variants significantly associated with usage of a specific statin (Table 2). Variants were identified with usage of simvastatin, atorvastatin, and fluvastatin, and no significant hits were identified for either rosuvastatin or pravastatin. Each of the significant hits from simvastatin and atorvastatin were significant in the hypercholesterolemia GWAS, while the fluvastatin significant variant was not. Two of the significantly associated variants have been previously associated with statin response: rs10455872 in lipoprotein(a) (*LPA*) has been associated with statin efficacy and toxicity^26,27^, and rs429358 in Apolipoprotein E (*APOE*) has been associated with simvastatin efficacy^28^. The variant found to be significantly associated with fluvastatin usage is intergenic and not located near any known genes or functional regions. Manhattan plots for the DS-GWAS with significant variants associations are shown in Figure 2. Both variants associated with atorvastatin selection were also significantly associated with simvastatin, although with the odds ratio in the opposite direction. The variants identified as significant in simvastatin (rs10455872 & rs74617384) and atorvastatin in *LPA* and *APOE* (rs429358) were reported to be significantly associated with cholesterol and other lipid levels in UK Biobank dataset displayed in Global Biobank Engine. The variant associated with fluvastatin (rs10492923) has not been previously associated with any phenotype.

**Table 2.**
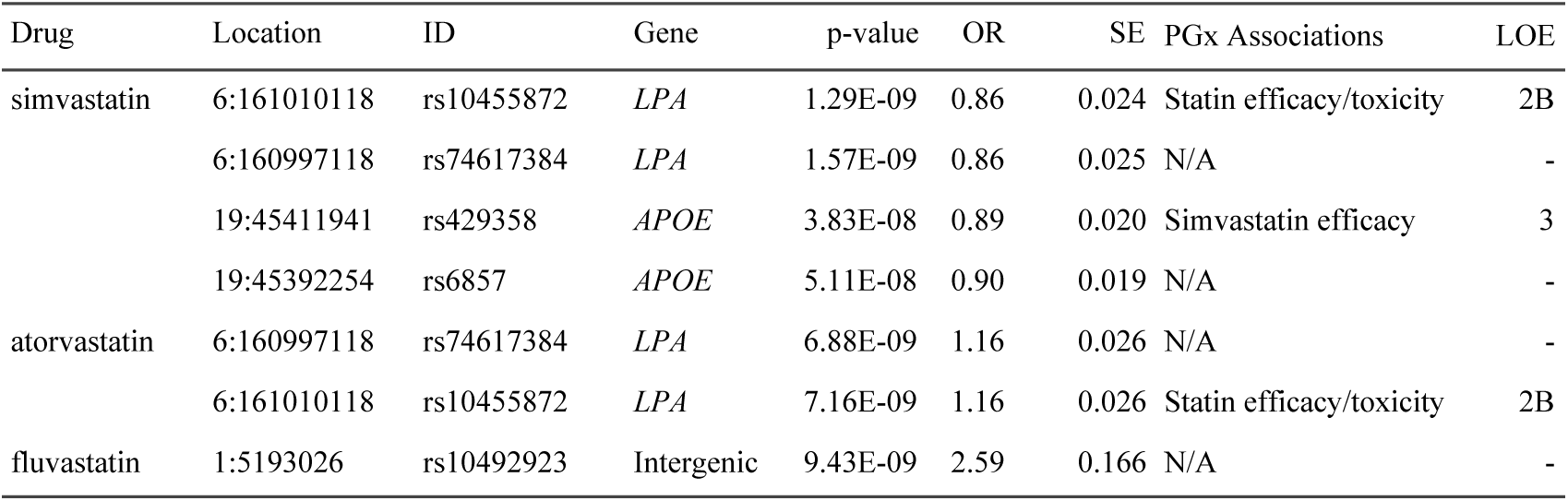
DS-GWAS significant association for statin selection. LOE indicates the level of evidence for the existing drug-variant relationship in PharmGKB. An LOE of 2B indicates there is moderate evidence for the variant-drug association, while an LOE of 3 indicates a low level of existing evidence.

| Drug | Location | ID | Gene | p-value | OR | SE | PGx Associations | LOE |
| --- | --- | --- | --- | --- | --- | --- | --- | --- |
| simvastatin | 6:161010118 | rs10455872 | *LPA* | 1.29E-09 | 0.86 | 0.024 | Statin efficacy/toxicity | 2B |
|  | 6:160997118 | rs74617384 | *LPA* | 1.57E-09 | 0.86 | 0.025 | N/A | - |
|  | 19:45411941 | rs429358 | *APOE* | 3.83E-08 | 0.89 | 0.020 | Simvastatin efficacy | 3 |
|  | 19:45392254 | rs6857 | *APOE* | 5.11E-08 | 0.90 | 0.019 | N/A | - |
| atorvastatin | 6:160997118 | rs74617384 | *LPA* | 6.88E-09 | 1.16 | 0.026 | N/A | - |
|  | 6:161010118 | rs10455872 | *LPA* | 7.16E-09 | 1.16 | 0.026 | Statin efficacy/toxicity | 2B |
| fluvastatin | 1:5193026 | rs10492923 | Intergenic | 9.43E-09 | 2.59 | 0.166 | N/A | - |

**Figure 2.**
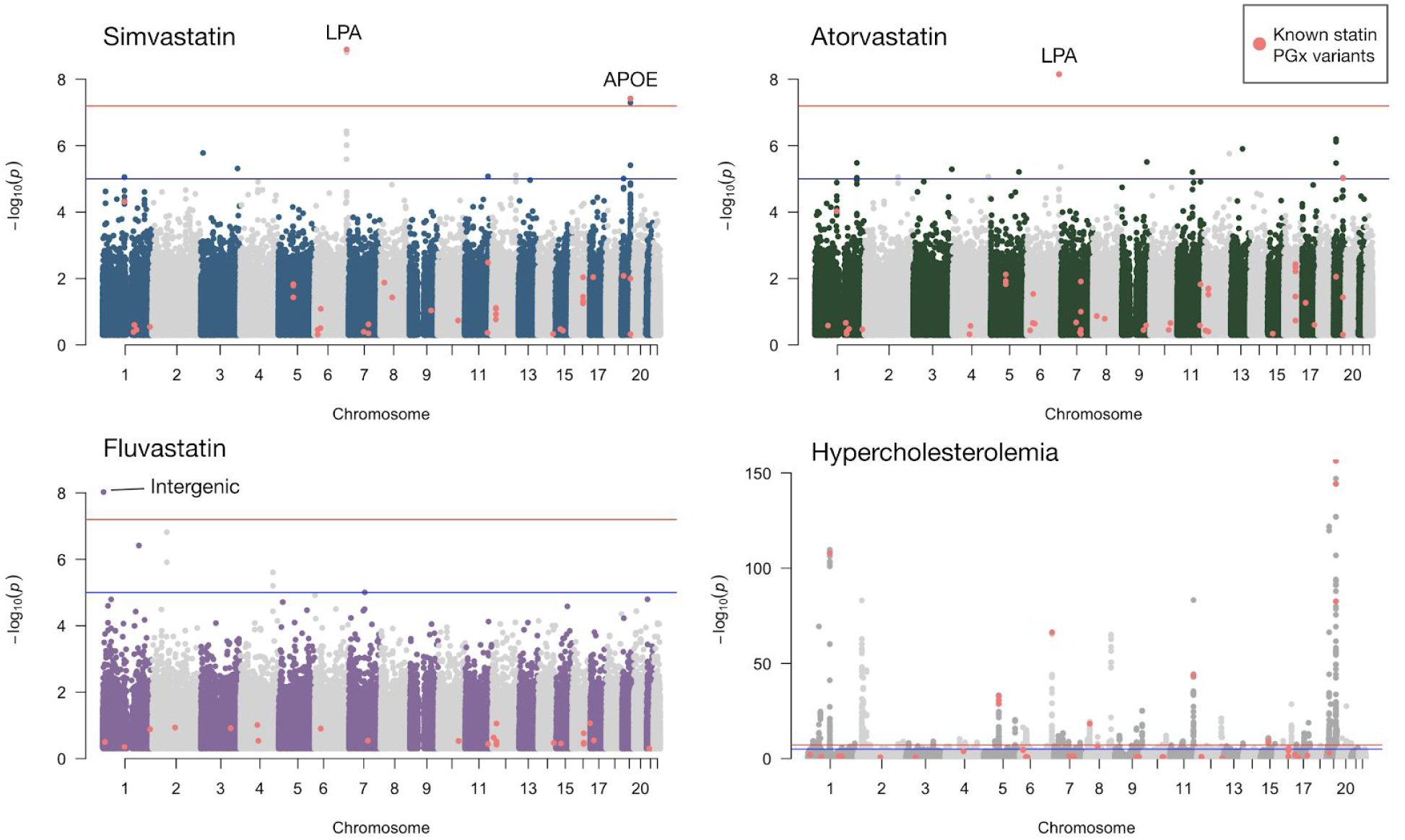
Manhattan plots for DS-GWAS of individual statins. Variants highlighted in green were found to be significantly associated with hypercholesterolemia. GWAS of simvastatin users compared to all other statin users recovers known pharmacogenetic variants for statin response in *LPA* and *APOE*. Atorvastatin DS-GWAS recovers two LPA variants previously reported for association to statin response. DS-GWAS for fluvastatin users identifies one potential statin response variant in an intergenic region on chromosome 1. We also show a Manhattan plot for a hypercholesterolemia GWAS for comparison. Pink points represent known statin pharmacogenetic variants.

We calculated the pearson correlation coefficient between z-scores from a hypercholesterolemia GWAS and three GWAS, a GWAS comparing all statin users and all non-statin users, and DS-GWAS for atorvastatin and simvastatin. Hypercholesterolemia and statin users had a correlation coefficient of 0.99, atorvastatin and hypercholesterolemia 0.80, and simvastatin and hypercholesterolemia −0.80 (Supplemental Figure 1). All z-score distributions were significantly correlated, (p ∼ 0).

### Genotype association with drug dose

In addition to testing association with drug selection, we investigated whether variants significantly associated with drug choice were also associated with dosage. Using the primary care data we calculated the prescribed daily dose for subjects on simvastatin, atorvastatin, rosuvastatin, and fluvastatin. Using the NICE guidelines, we then binned individuals based on the intensity of the statin regimen. We selected a representative variant for each peak to use for the analysis, three variants total. We found that two of the three variants significantly associated with statin selection were also associated with atorvastatin dose (rs429358: OR = 1.7, *P* = 1.6× 10^−4^, p(high intensity) = 0.163, p(not high intensity = 0.143), rs74617384: OR = 1.4, *P* = 5.2 × 10^−3^, p(high intensity) = 0.166, p(not high intensity = 0.149)). Dose associations for rs429358 and rs74617384 from *APOE* and *LPA*, respectively, are shown in Figure 3. These variants were only significantly associated with atorvastatin regimen, no other drugs. The chromosome 1 variant identified in the fluvastatin DS-GWAS was not associated with the dosing regimen for any drug.

**Figure 3.**
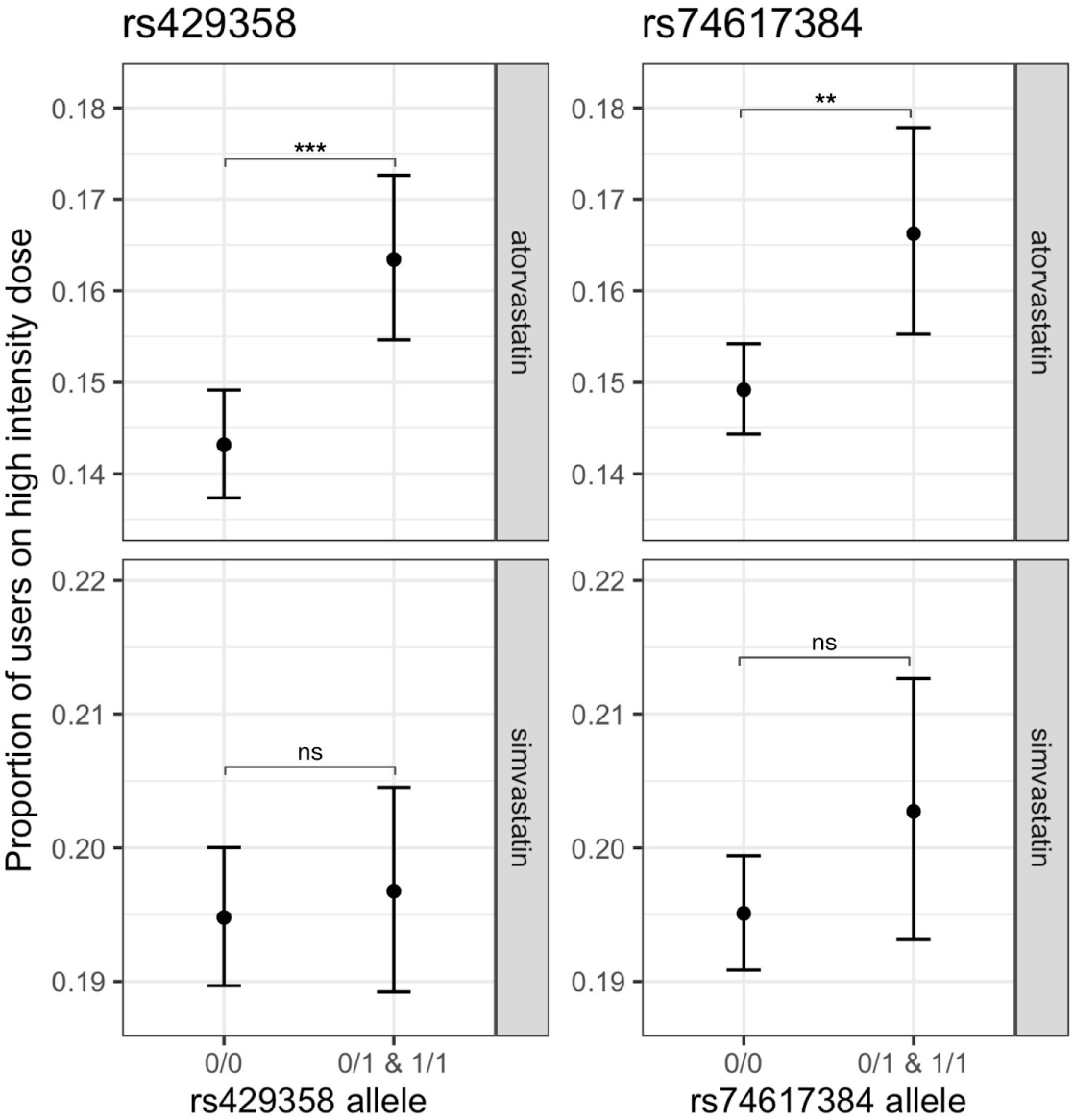
Variant association with high intensity statin regimen. The proportion of users of a high intensity dose regimen is shown here for two carriers of two variants on two statins, simvastatin and atorvastatin. Both variants are significantly associated with a high intensity regimen among individuals who carry an alternate allele. A high intensity is defined as a prescribed daily dose greater than or equal to 40mg of atorvastatin or 80mg of simvastatin. Both variants were significantly associated with a decreased likelihood of being on simvastatin (a weaker statin) and rs74617384 was significantly associated with an increased likelihood of being on atorvastatin (a stronger statin).

### Known statin pharmacogenetic analysis

Finally, we sought to evaluate known statin pharmacogenetics in the UK Biobank using the two methods previously described, GWAS and dose association. We queried 104 variants from PharmGKB reported to be associated with statin efficacy or toxicity with any level of evidence. Of those, 22 were found to be associated with treatment intensity with dose intensity or statin selection (Table 3). The two most significant variants associated with dose were significant in the simvastatin DS-GWAS, rs10455872 in *LPA* and rs429358 in *APOE*. In all, fourteen unique genes are found to have a significant association with dosage in at least one statin. Atorvastatin intensity is the most frequently associated with PGx variants (15 associations), but each drug is associated with at least one variant. Dosage plots for all variants included in Supplemental File 2.

**Table 3.**
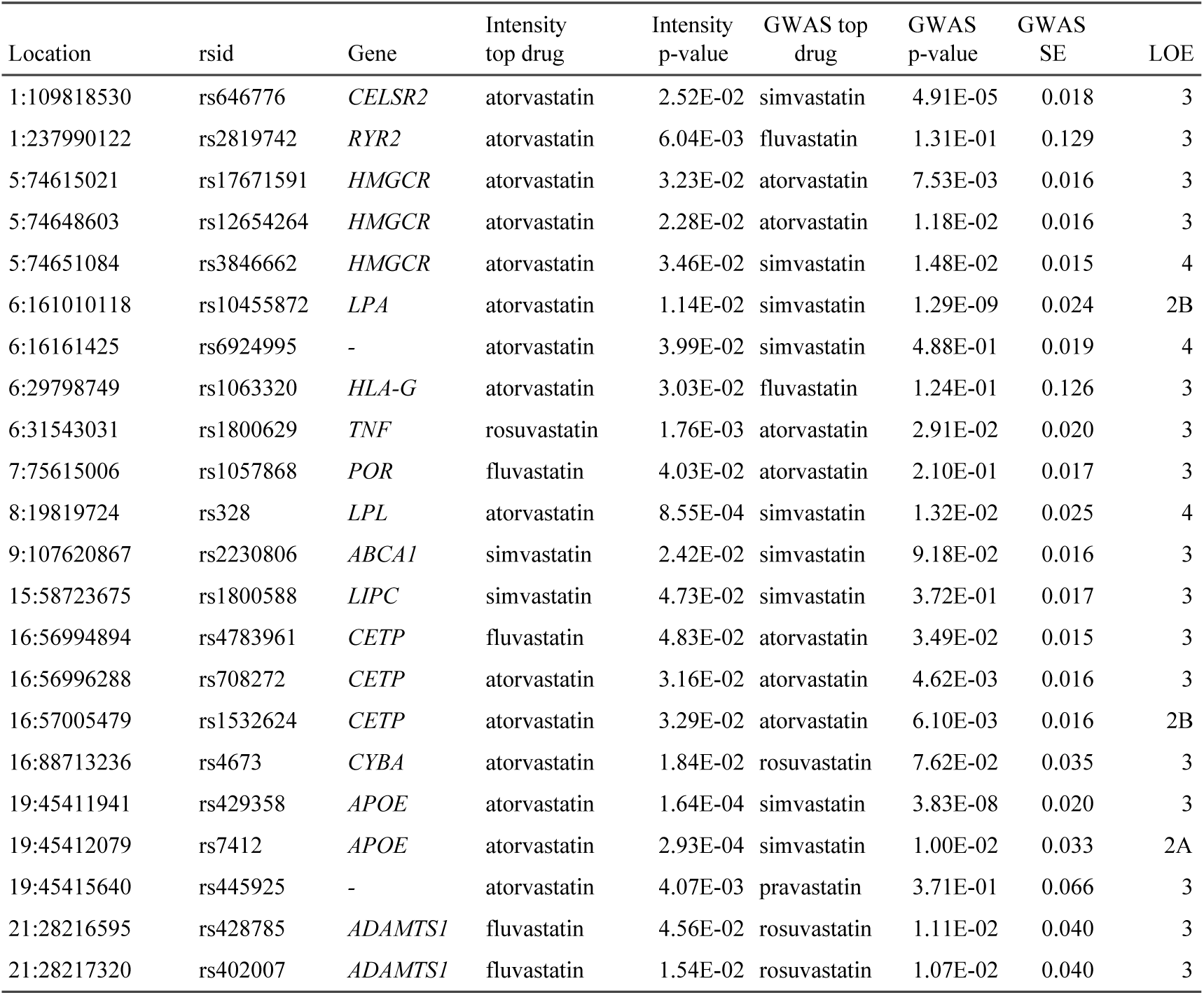
Known pharmacogenetic associations with statins. We determined the most significant association between statin-associated variants in PharmGKB from DS-GWAS and higher intensity regimens. The drug with the most significant association is shown in for each comparison is shown in “Intensity top drug” and “GWAS top drug”. LOE represents the minimum level of evidence for each variant in PharmGKB. This table is sorted by genomic location.

## Discussion

In this study we use DS-GWAS to identify genetic variants that may lead an individual to take one type of statin over another, and determine whether these variants also lead individuals to take different doses of statins as a result of routine medical treatment. We identify several variants in *LPA* and *APOE* significantly associated with a decreased propensity for taking a moderate statin (simvastatin), and that the *LPA* variants are associated with taking a stronger statin (atorvastatin). Further, we show that individuals who carry an alternate allele for any of these variants are significantly more likely to be on a high intensity dosing regimen once on atorvastatin.

Variants within *LPA* and *APOE* have previously been associated with statin efficacy^29–32^. Previous studies have shown that carriers of the *LPA* variant rs10455872 have a decreased response to statins. LDL-c levels are reduced by 5.9% less among carriers per alternate allele than among non-carriers^30^. This supports the hypothesis that carriers of this variant will require a stronger therapeutic intervention to achieve the desired effect, which this work shows often results in using a stronger statin and a higher dose of the stronger statin. Further, drug-resistant hypercholesterolemia has been linked to high levels of LPA, and individuals with high LPA levels are candidates for more intensive therapy^33^. The intronic *LPA* variant rs10455872 identified in this study is known to increase LPA levels^34^. Variants in *APOE*, including rs429358, have been linked to significant increase in LDL-c, but also a significant reduction in LDL-c levels in response to statin therapy^35^. Previous studies of rs429358 influence on statin response, however, are mixed; some studies find that LDL-c levels in carriers do not decrease as much as in non-carriers^36,37^. Our results show that carriers of rs429358 are more likely to be on a stronger statin and a higher dose of the stronger statin.

We found evidence to support 22 variants in 14 genes associated with statin response in PharmGKB in the form of an association with dosage. Each of these variants is significantly associated with carriers being on a different statin intensity than those who are homozygous reference. This additional evidence may encourage further study and clinical validation.

The comparison of z-scores between the hypercholesterolemia GWAS and the DS-GWAS for atorvastatin and simvastatin show that there is a significant association between directionality of effect for a variants effect on high cholesterol risk and statin selection. Variants with a larger effect size for high cholesterol tend to be correlated with being on a strong statin, such as atorvastatin, and have an opposite effect for the selection of simvastatin. However, we find that the variants that are most significantly associated with statin selection (in *LPA*) were not the most significantly associated with high cholesterol. The identification of these genomic variants that have been previously reported to be associated with statin response represent the potential of using DS-GWAS a method of identifying variants associated with drug response.

Fluvastatin had a single novel association identified through DS-GWAS. The fluvastatin hit was found in a non-coding region on chr1, at 5193026 bp. We found no known previous associations to these variants or in the surrounding region. This novel variant associations with fluvastatin may warrant follow-up investigation given the validity of the *LPA* and *APOE* associations identified in the atorvastatin and simvastatin DS-GWAS.

Here we introduce the concept of a Drug Selection Genome-wide Association Study (DS-GWAS). The aim of this study design is to test whether there exists a genetic association between the prescribing of a particular drug and a particular genotype. For many diseases, the mechanism of selecting a particular drug to prescribe or dosing a selected drug still consists of trial-and-error^2,3^. Although adoption of PGx into clinical practice is increasing, there are still many drugs that lack PGx guidance and further research is needed^38,39^. We believe that biobanks that contain both genetic and prescribing data represent an opportunity to cheaply expand our understanding of PGx, but only if new study designs are created that enable us to leverage the information they contain^1^. The goal of the DS-GWAS is to capture the latent PGx signals created by clinicians during their trial-and-error process of prescribing medications to patients. We seek to identify genetic associations that potentially drive those decisions and this work serves as a proof of concept that this is possible. As with any GWAS, the challenge lies with biological interpretation of the variants identified. As variants may be related to any number of reasons for choosing a particular drug, thorough validation of significant variants will need to be carefully performed for those variants not previously identified to have any meaning.

## Contributions

A.L., G.M. and R.A. conceived the idea for the project. A.L. and G.M. executed the cohort building, performed the genome-wide association studies and conducted the analysis/interpretation of the genome-wide association study summary statistics. Y.T. performed quality control of the genotyping data. M.A.R. oversaw the project and aided statistical analyses.

## Acknowledgements

A.L. is supported by the National Science Foundation DGE 1656518. G.M. is supported by the Big Data to Knowledge (BD2K) from the National Institutes of Health (T32 LM012409). Y.T. is supported by Funai Overseas Scholarship from Funai Foundation for Information Technology and the Stanford University School of Medicine. R.B.A is supported by NIH/National Institute of General Medical Sciences PharmGKB resource (U24HG010615), NIH GM102365, and the Chan Zuckerberg Biohub. This research has been conducted using the UK Biobank Resource under Application Numbers 24983 and 33722. We thank all the participants in the UK Biobank study. Most of the computing for this project was performed on the Sherlock cluster. We would like to thank Stanford University, the PharmGKB resource (NIH HG010615), and the Stanford Research Computing Center for providing the computational resources that contributed to these research results. We would also like to thank Christopher Chang for his work optimizing Plink2.

## Competing Interests

R.B.A. is a stockholder in Personalis.com, 23andme.com. M.A.R. is on the SAB of 54Gene and Computational Advisory Board for Goldfinch Bio and has advised BioMarin, Third Rock Ventures, MazeTx and Related Sciences.

