## Supplemental File 1 for "*LPA* and *APOE* are associated with statin selection in the UK Biobank"

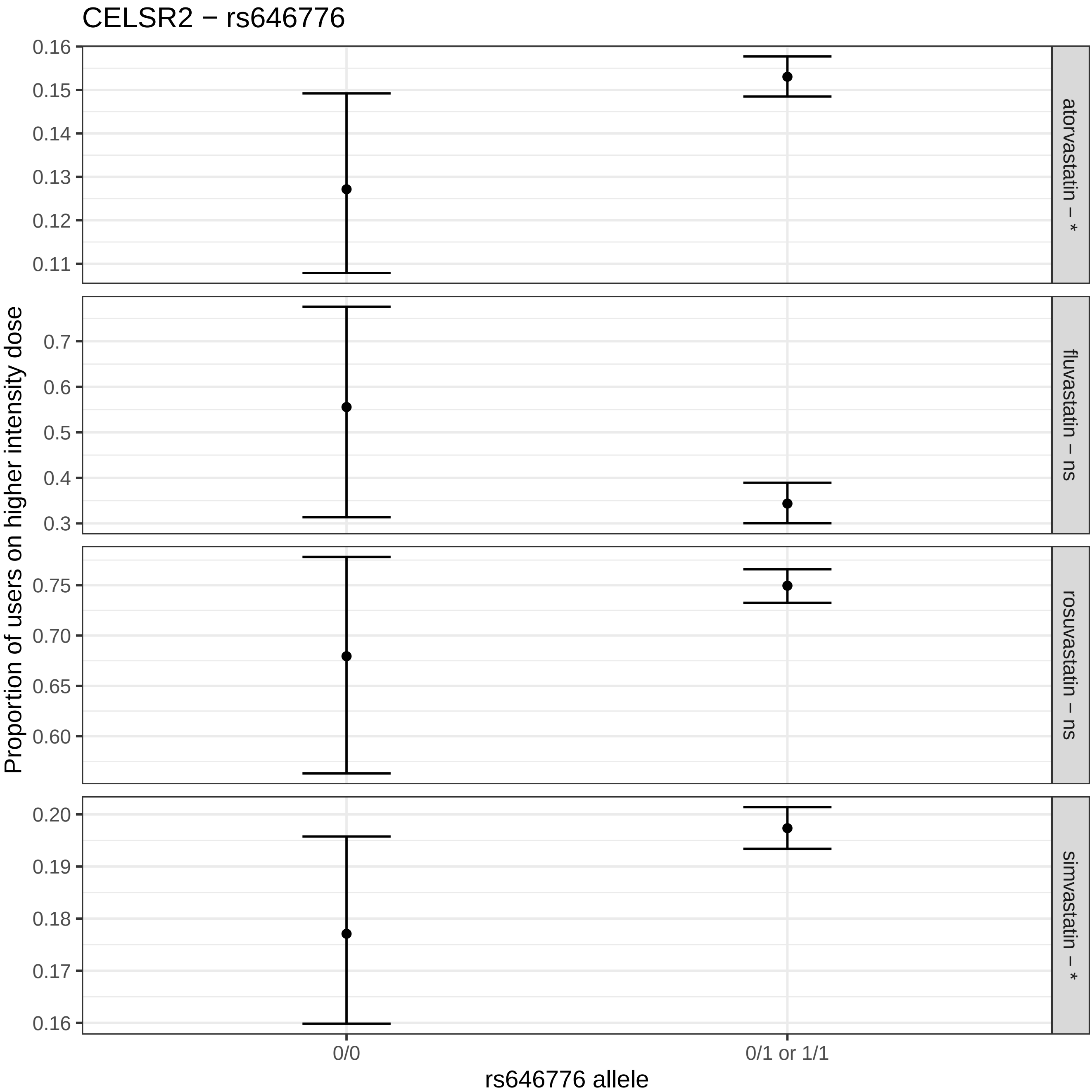
CELSR2 − rs646776
Proportion of users on higher intensity dose
atorvastatin – *
fluvastatin – ns
rosuvastatin – ns
simvastatin – *
0.16
0.15
0.14
0.13
0.12
0.11
0.7
0.6
0.5
0.4
0.3
0.75
0.70
0.65
0.60
0.20
0.19
0.18
0.17
0.16
0/0
0/1 or 1/1
rs646776 allele

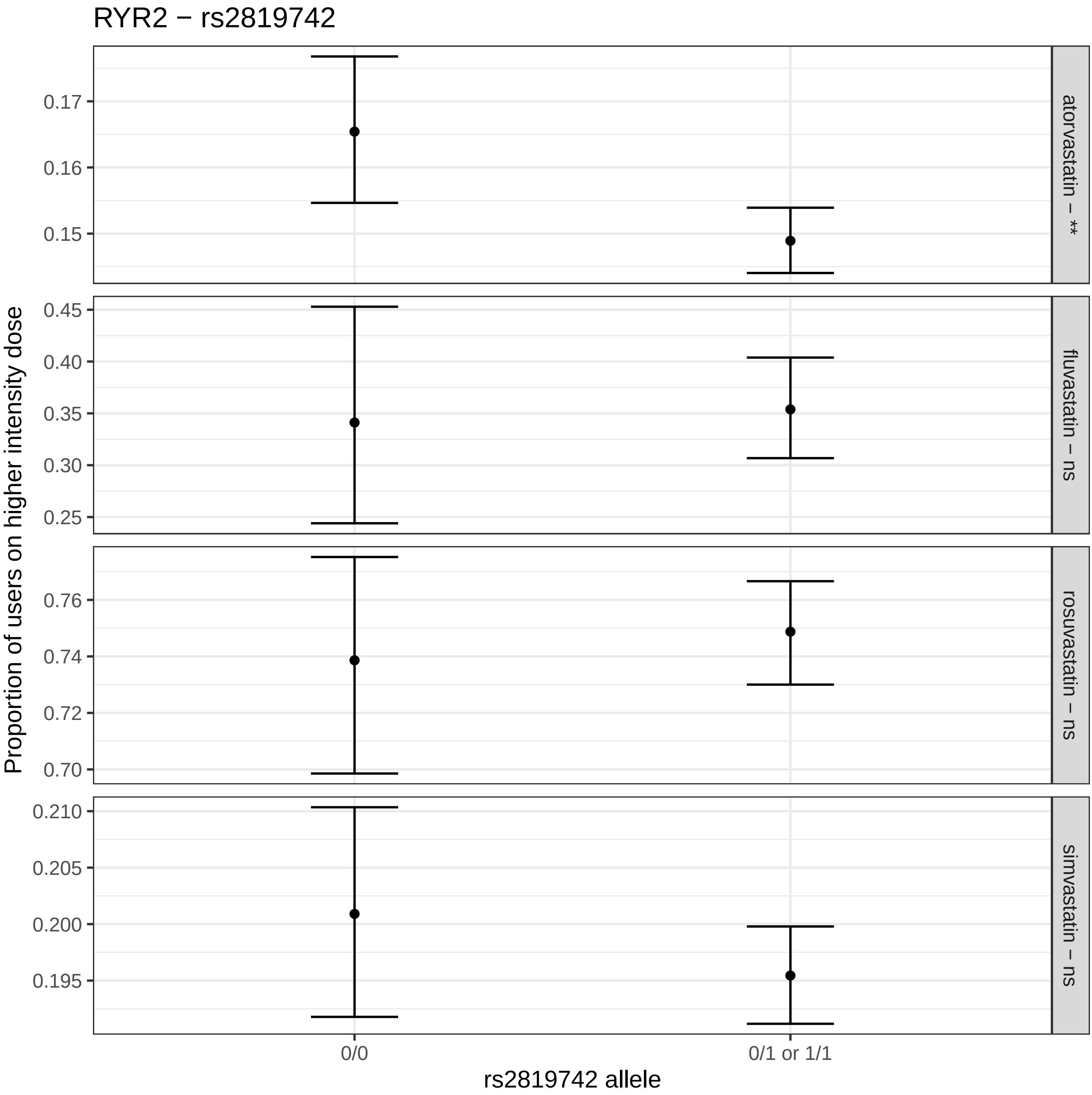

RYR2 − rs2819742
Proportion of users on higher intensity dose
rs2819742 allele
0/0
0/1 or 1/1
atorvastatin − **
fluvastatin − ns
rosuvastatin − ns
simvastatin − ns
0.17
0.16
0.15
0.45
0.40
0.35
0.30
0.25
0.76
0.74
0.72
0.70
0.210
0.205
0.200
0.195

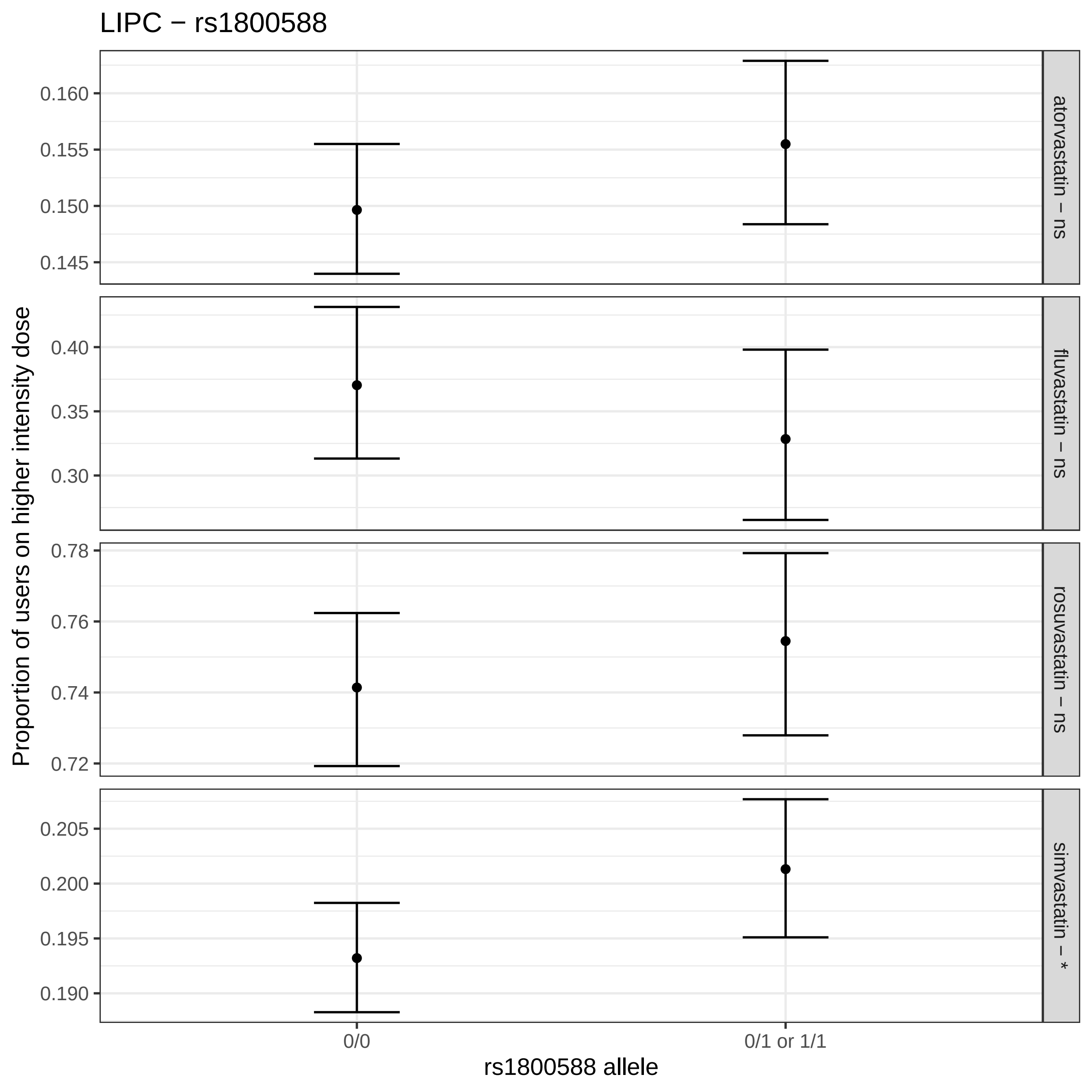

LIPC − rs1800588
Proportion of users on higher intensity dose
0.160
0.155
0.150
0.145
atorvastatin − ns
0.40
0.35
0.30
fluvastatin − ns
0.78
0.76
0.74
0.72
rosuvastatin − ns
0.205
0.200
0.195
0.190
simvastatin − *
0/0
0/1 or 1/1
rs1800588 allele

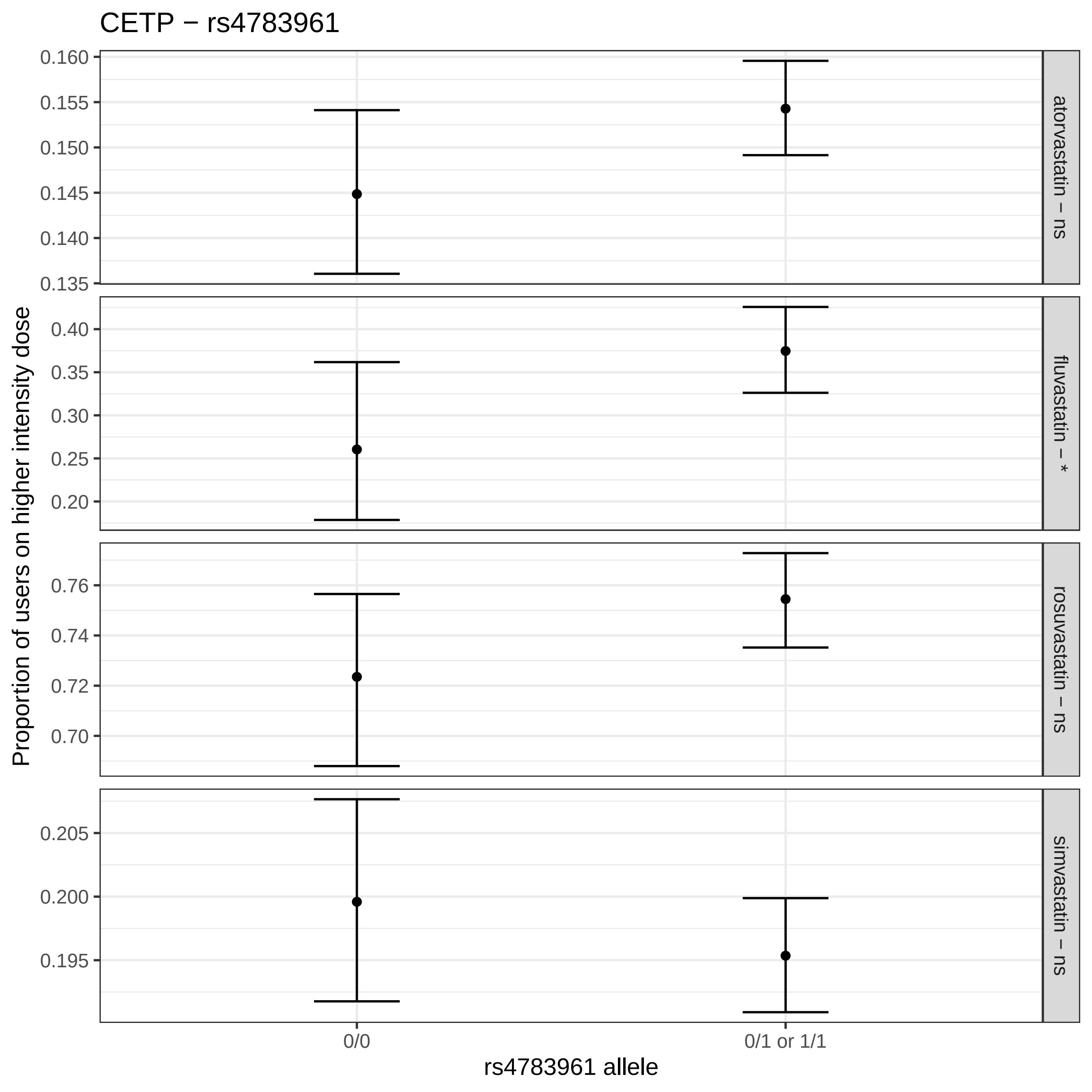

CETP − rs4783961
Proportion of users on higher intensity dose
atorvastatin − ns
fluvastatin − *
rosuvastatin − ns
simvastatin − ns
0.160
0.155
0.150
0.145
0.140
0.135
0.40
0.35
0.30
0.25
0.20
0.76
0.74
0.72
0.70
0.205
0.200
0.195
0/0
0/1 or 1/1
rs4783961 allele

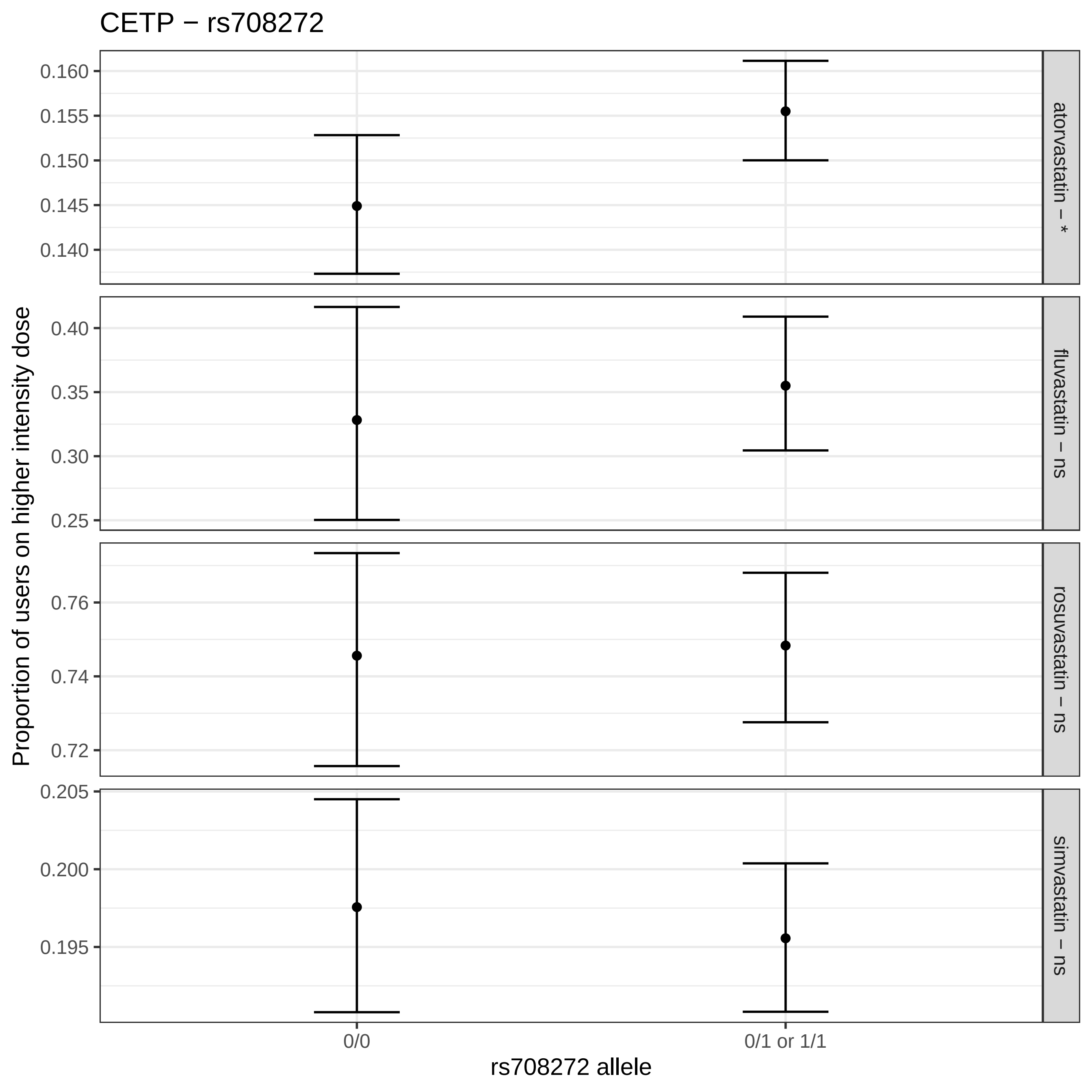
CETP − rs708272
0.160
0.155
0.150
0.145
0.140
atorvastatin − *
0.40
0.35
0.30
0.25
fluvastatin − ns
0.76
0.74
0.72
rosuvastatin − ns
0.205
0.200
0.195
simvastatin − ns
Proportion of users on higher intensity dose
0/0
0/1 or 1/1
rs708272 allele

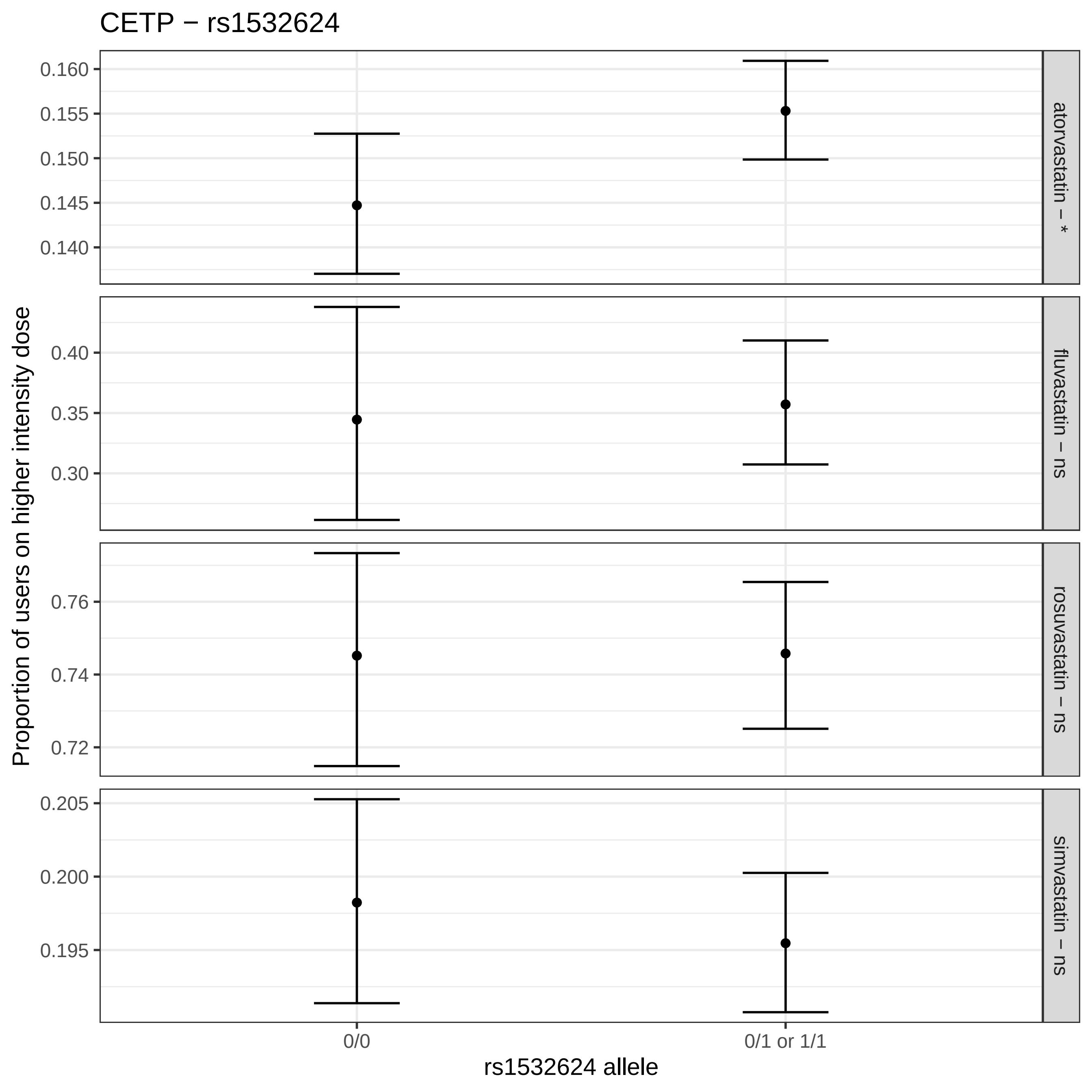

CETP − rs1532624
Proportion of users on higher intensity dose
atorvastatin − *
fluvastatin − ns
rosuvastatin − ns
simvastatin − ns
0.160
0.155
0.150
0.145
0.140
0.40
0.35
0.30
0.76
0.74
0.72
0.205
0.200
0.195
0/0
0/1 or 1/1
rs1532624 allele

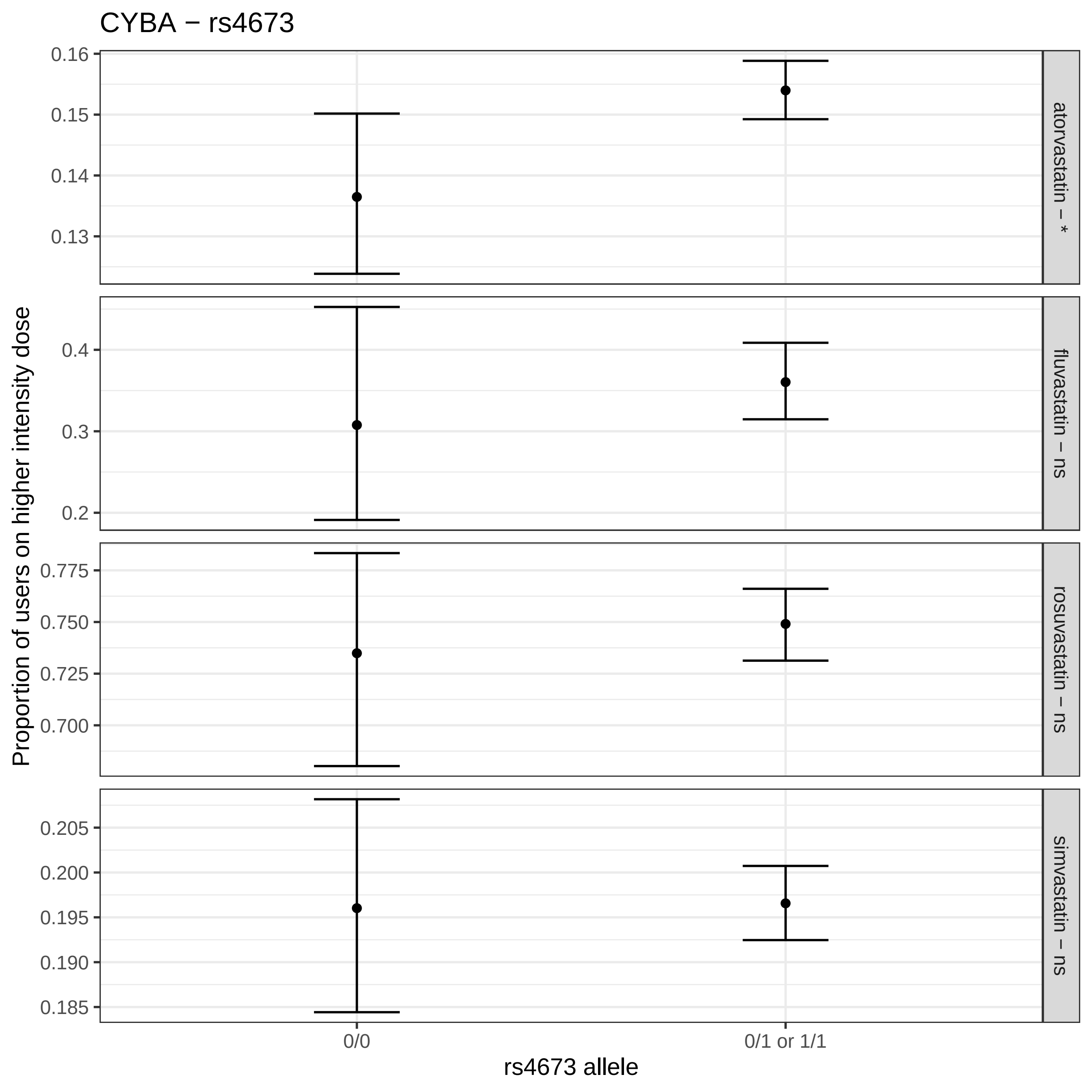

CYBA − rs4673
Proportion of users on higher intensity dose
0.160
0.150
0.140
0.130
0.4
0.3
0.2
0.775
0.750
0.725
0.700
0.205
0.200
0.195
0.190
0.185
atorvastatin − *
fluvastatin − ns
rosuvastatin − ns
simvastatin − ns
0/0
0/1 or 1/1
rs4673 allele

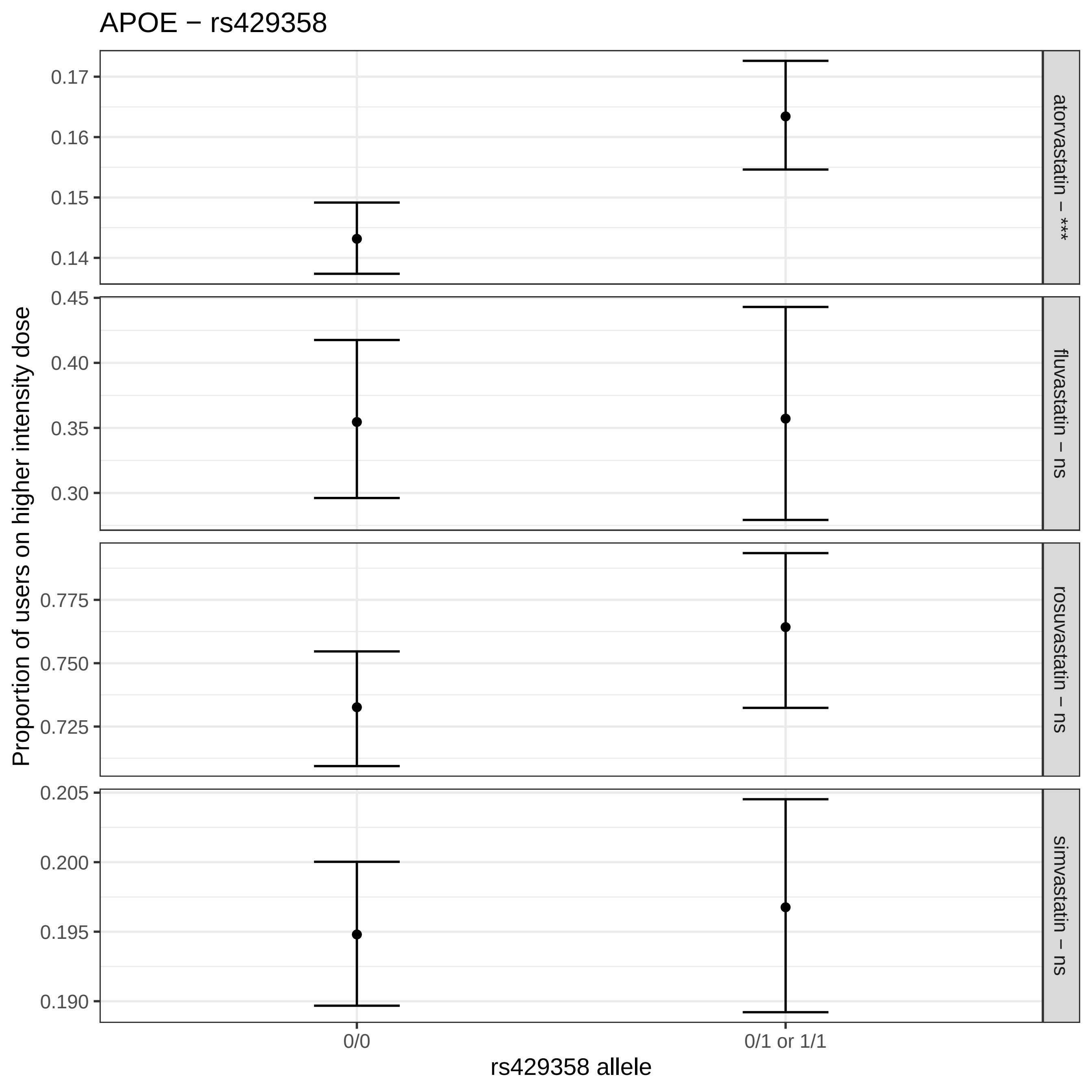
APOE − rs429358
Proportion of users on higher intensity dose
rs429358 allele
0/0
0/1 or 1/1
atorvastatin − ***
fluvastatin − ns
rosuvastatin − ns
simvastatin − ns
0.17
0.16
0.15
0.14
0.45
0.40
0.35
0.30
0.775
0.750
0.725
0.205
0.200
0.195
0.190

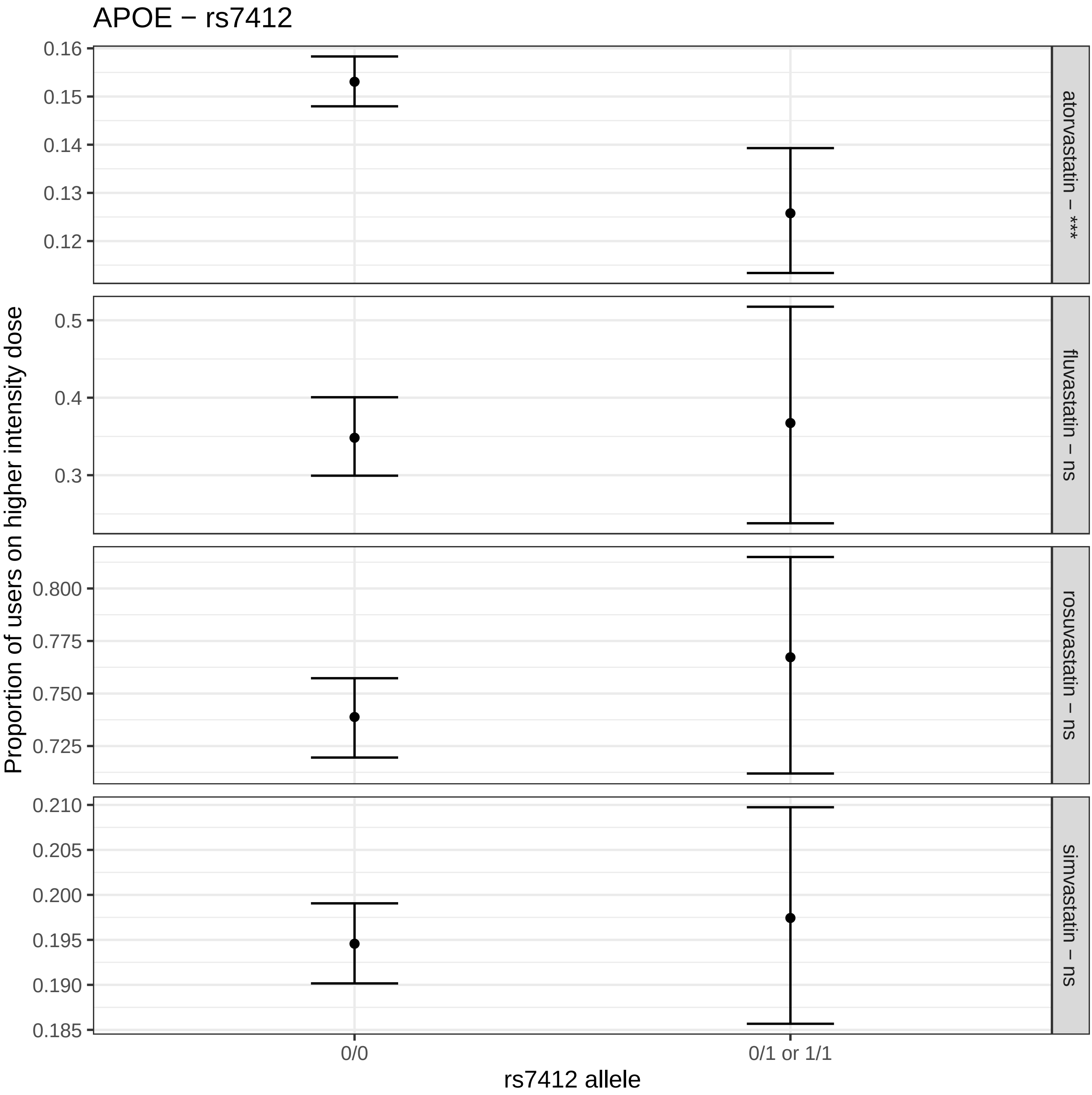

APOE − rs7412
Proportion of users on higher intensity dose
0.16
0.15
0.14
0.13
0.12
atorvastatin − ***
0.5
0.4
0.3
fluvastatin − ns
0.800
0.775
0.750
0.725
rosuvastatin − ns
0.210
0.205
0.200
0.195
0.190
0.185
simvastatin − ns
0/0
0/1 or 1/1
rs7412 allele

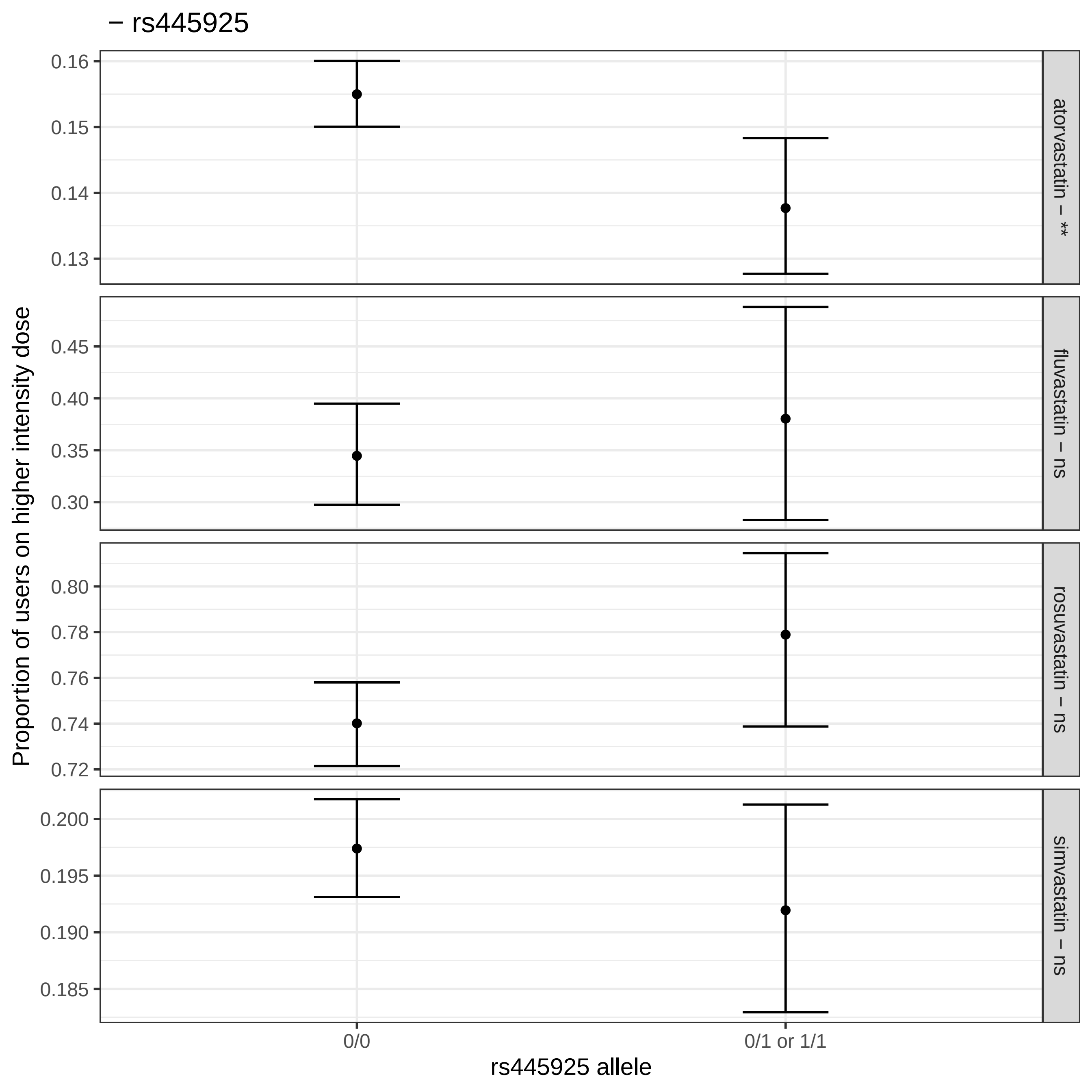

− rs445925
Proportion of users on higher intensity dose
rs445925 allele
0/0
0/1 or 1/1
atorvastatin – **
fluvastatin – ns
rosuvastatin – ns
simvastatin – ns
0.16
0.15
0.14
0.13
0.45
0.40
0.35
0.30
0.80
0.78
0.76
0.74
0.72
0.200
0.195
0.190
0.185

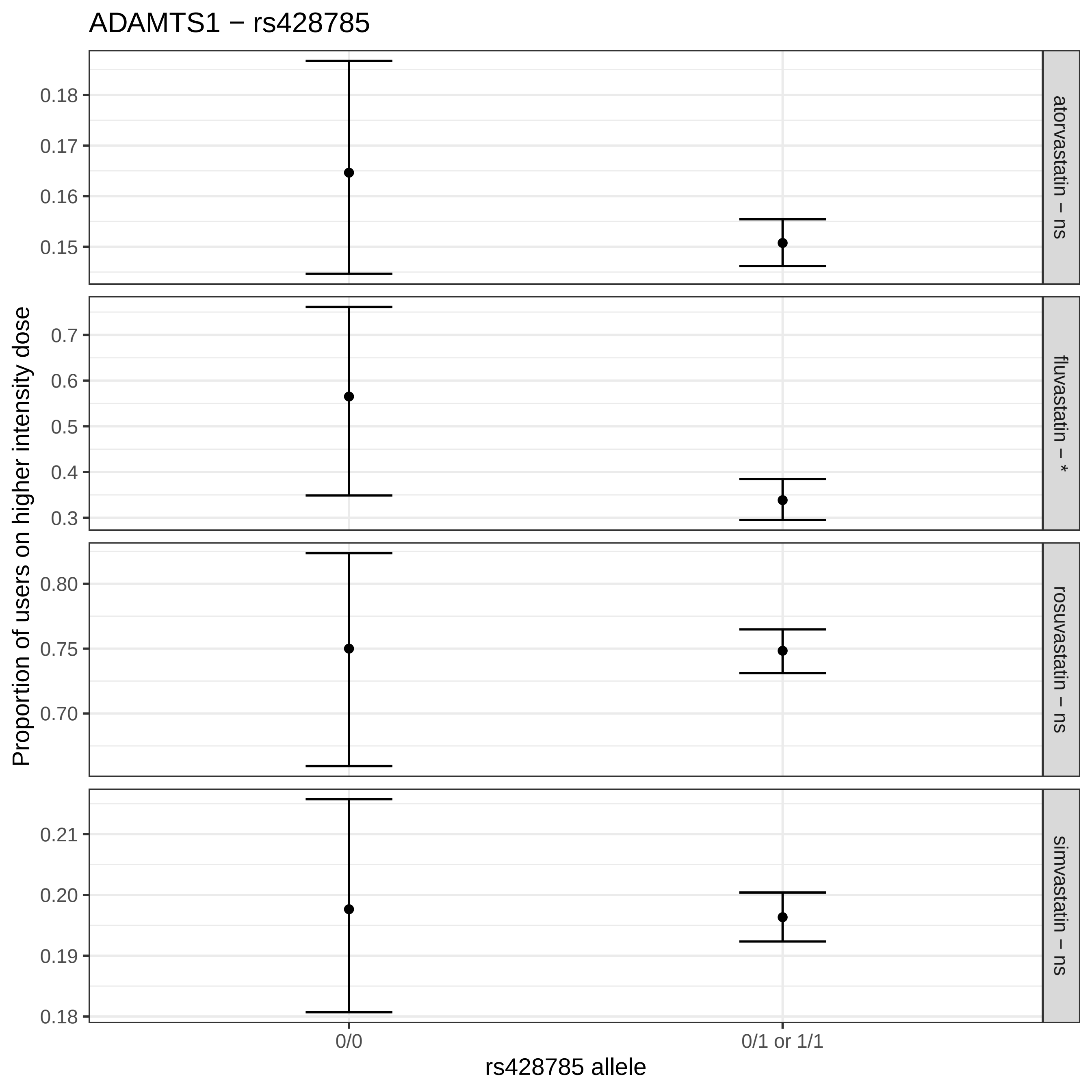
ADAMTS1 − rs428785
Proportion of users on higher intensity dose
atorvastatin – ns
fluvastatin – *
rosuvastatin – ns
simvastatin – ns
0.18
0.17
0.16
0.15
0.7
0.6
0.5
0.4
0.3
0.80
0.75
0.70
0.21
0.20
0.19
0.18
0/0
0/1 or 1/1
rs428785 allele

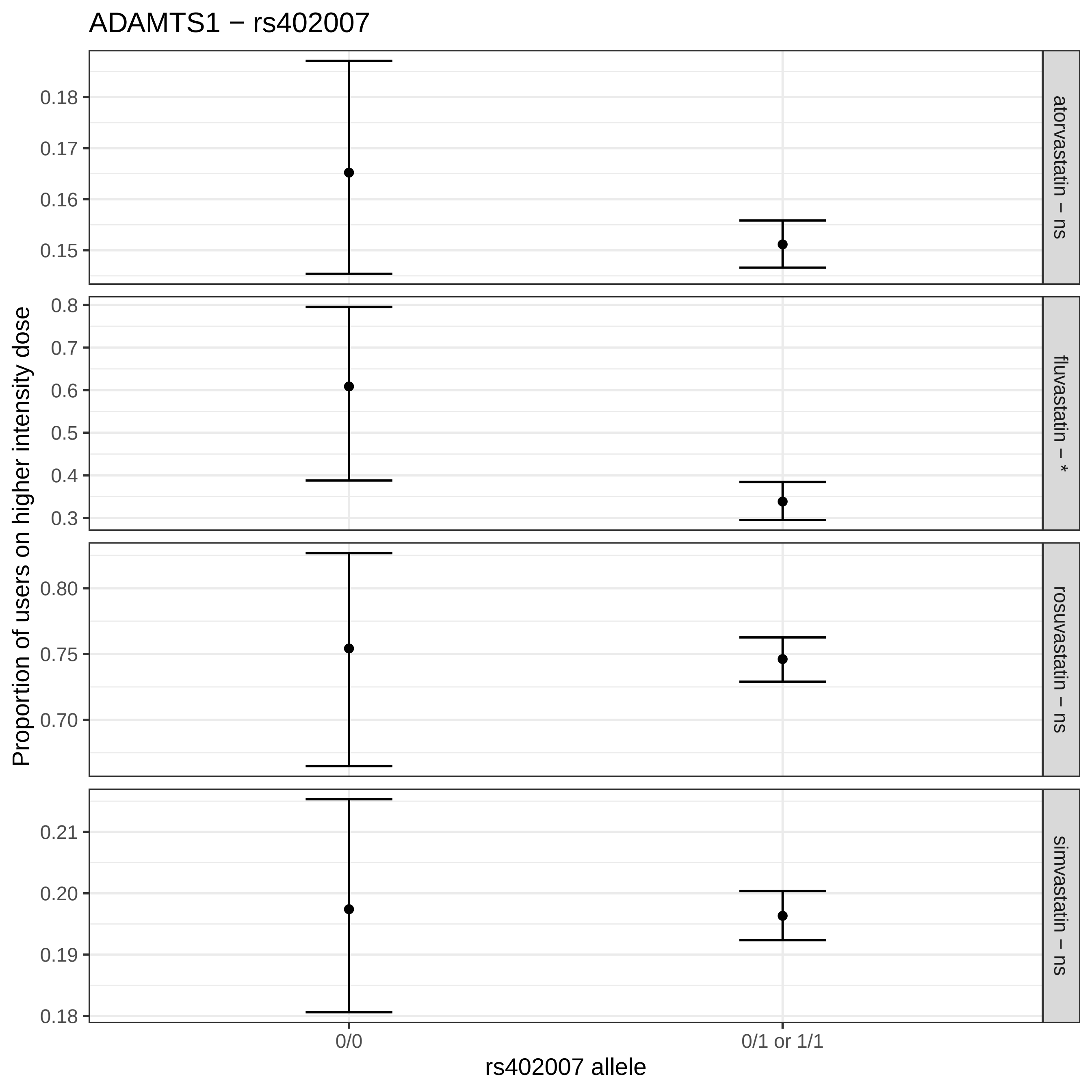
ADAMTS1 − rs402007
Proportion of users on higher intensity dose
atorvastatin − ns
fluvastatin − *
rosuvastatin − ns
simvastatin − ns
0.18
0.17
0.16
0.15
0.8
0.7
0.6
0.5
0.4
0.3
0.80
0.75
0.70
0.21
0.20
0.19
0.18
0/0
0/1 or 1/1
rs402007 allele

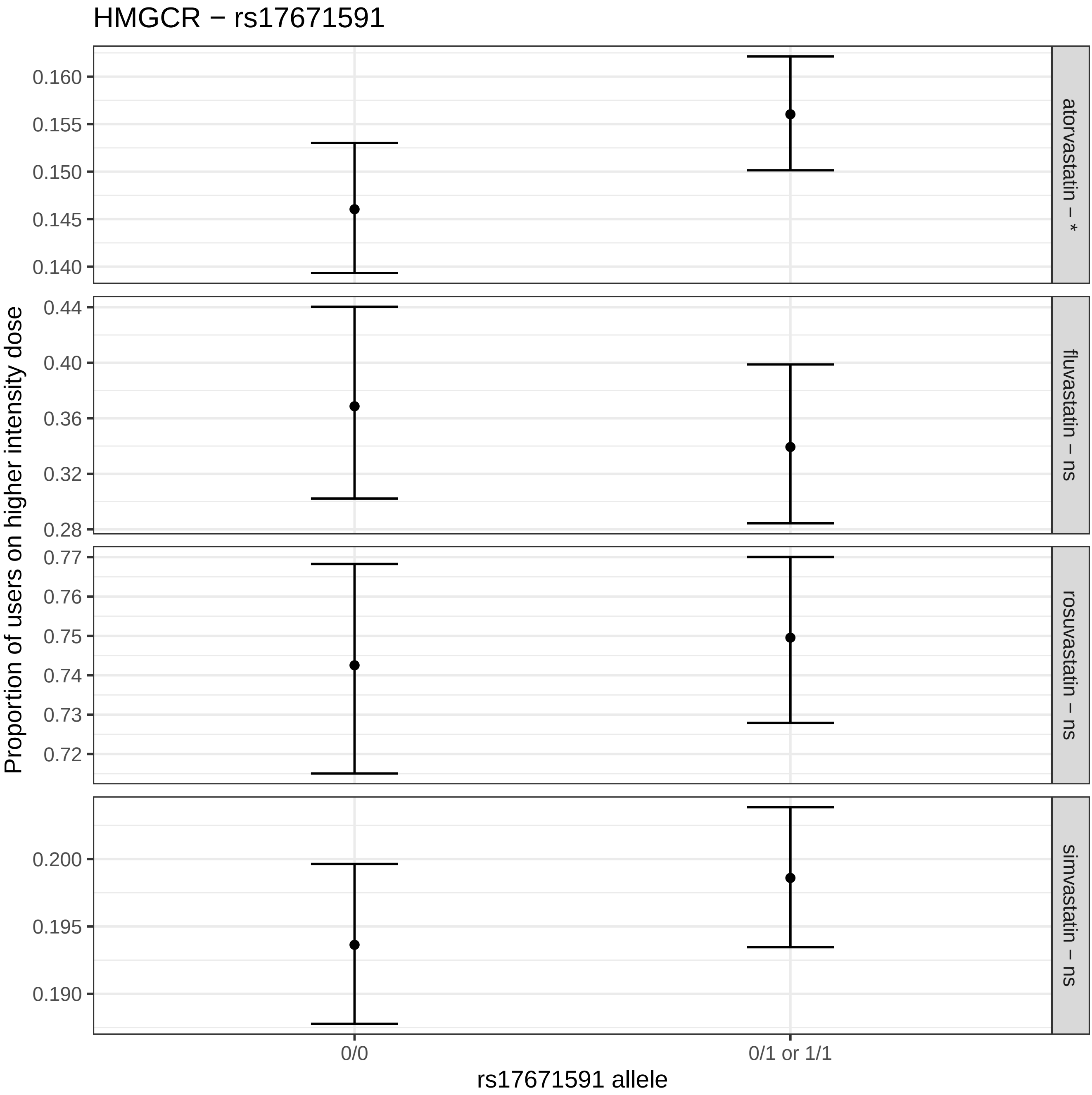
HMGCR − rs17671591
Proportion of users on higher intensity dose
atorvastatin − *
fluvastatin − ns
rosuvastatin − ns
simvastatin − ns
0.160
0.155
0.150
0.145
0.140
0.44
0.40
0.36
0.32
0.28
0.77
0.76
0.75
0.74
0.73
0.72
0.200
0.195
0.190
0/0
0/1 or 1/1
rs17671591 allele

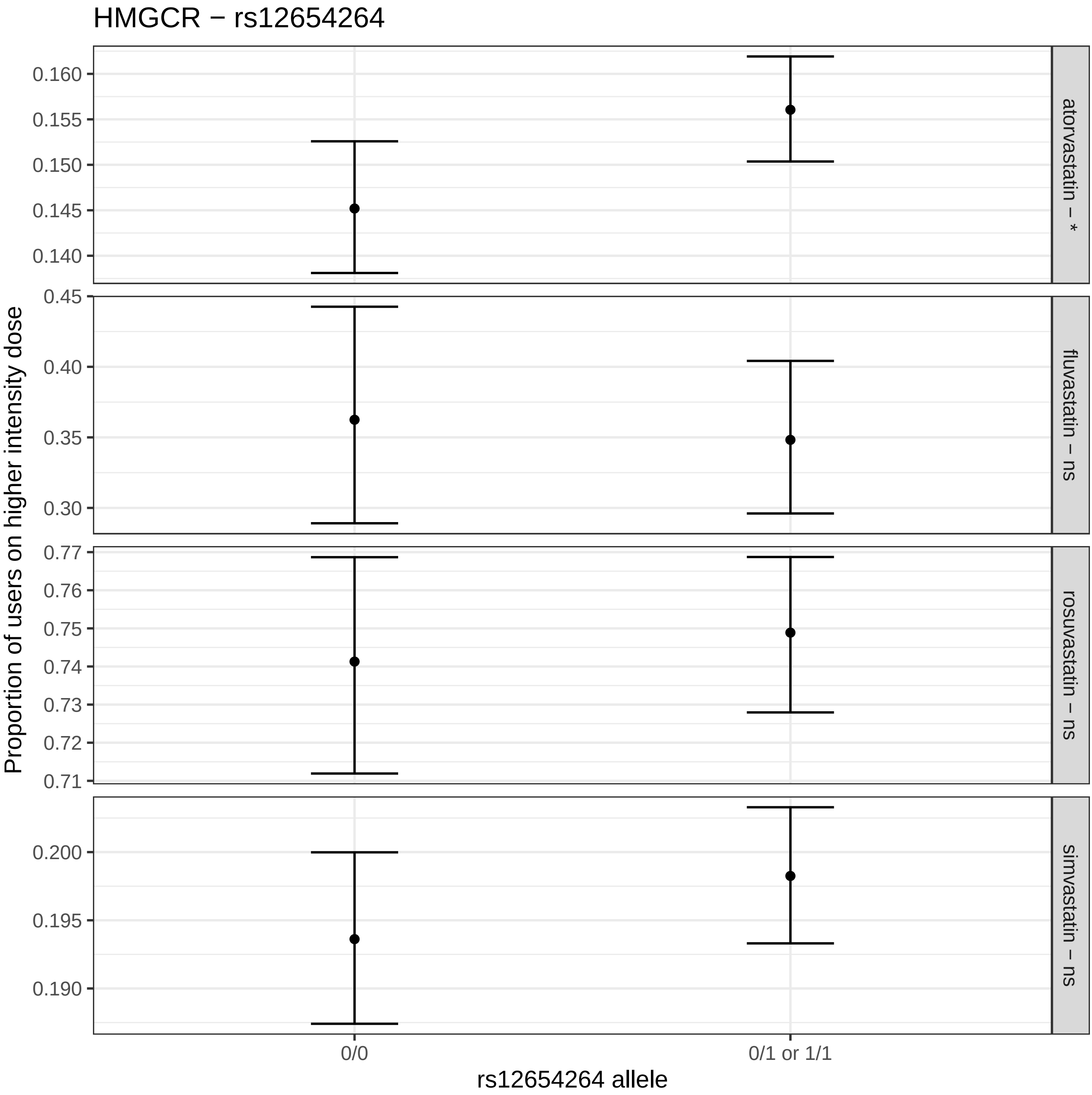

HMGCR − rs12654264
Proportion of users on higher intensity dose
0.160
0.155
0.150
0.145
0.140
0.45
0.40
0.35
0.30
0.77
0.76
0.75
0.74
0.73
0.72
0.71
0.200
0.195
0.190
atorvastatin − *
fluvastatin − ns
rosuvastatin − ns
simvastatin − ns
0/0
0/1 or 1/1
rs12654264 allele

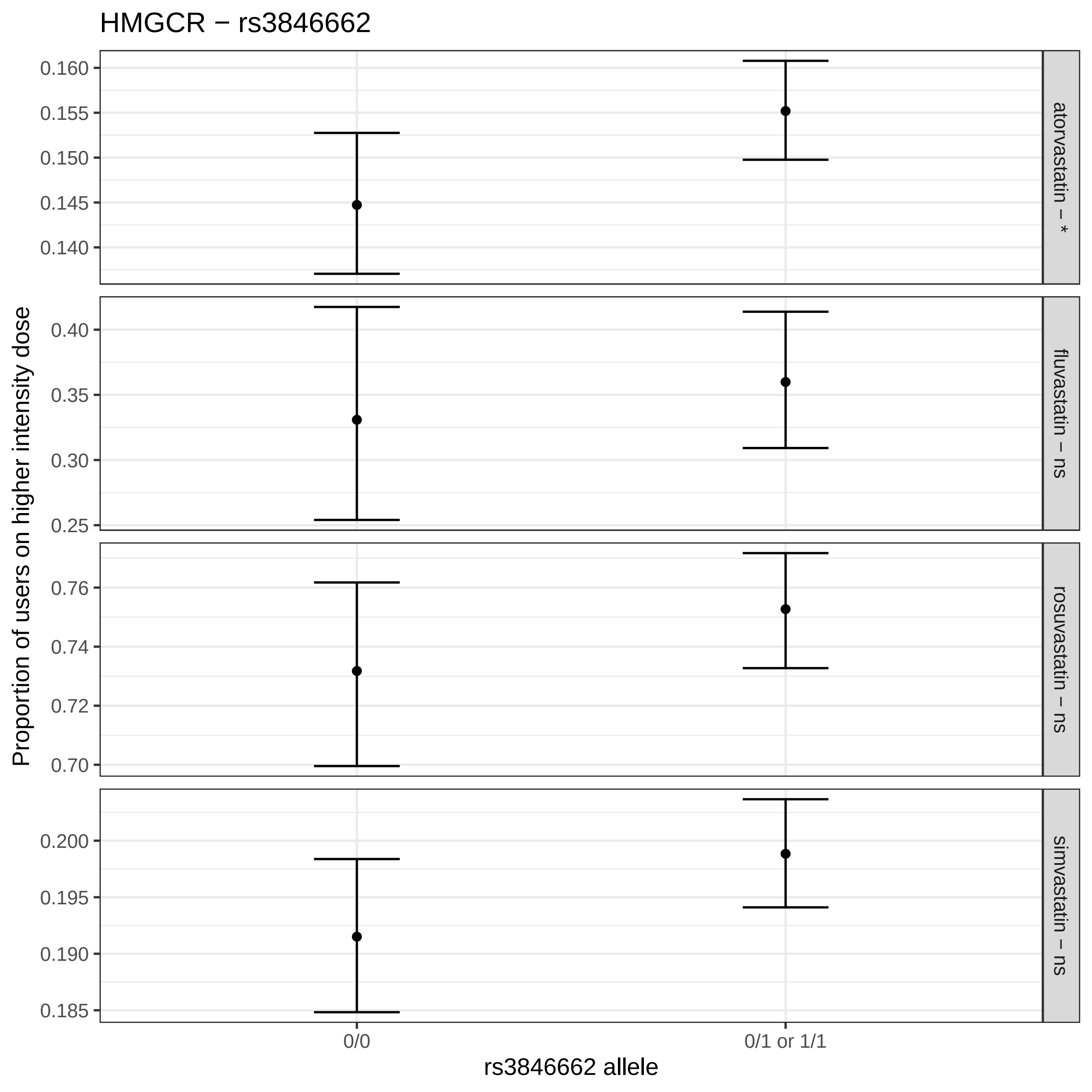
HMGCR − rs3846662
Proportion of users on higher intensity dose
atorvastatin – *
fluvastatin – ns
rosuvastatin – ns
simvastatin – ns
0.160
0.155
0.150
0.145
0.140
0.40
0.35
0.30
0.25
0.76
0.74
0.72
0.70
0.200
0.195
0.190
0.185
0/0
0/1 or 1/1
rs3846662 allele

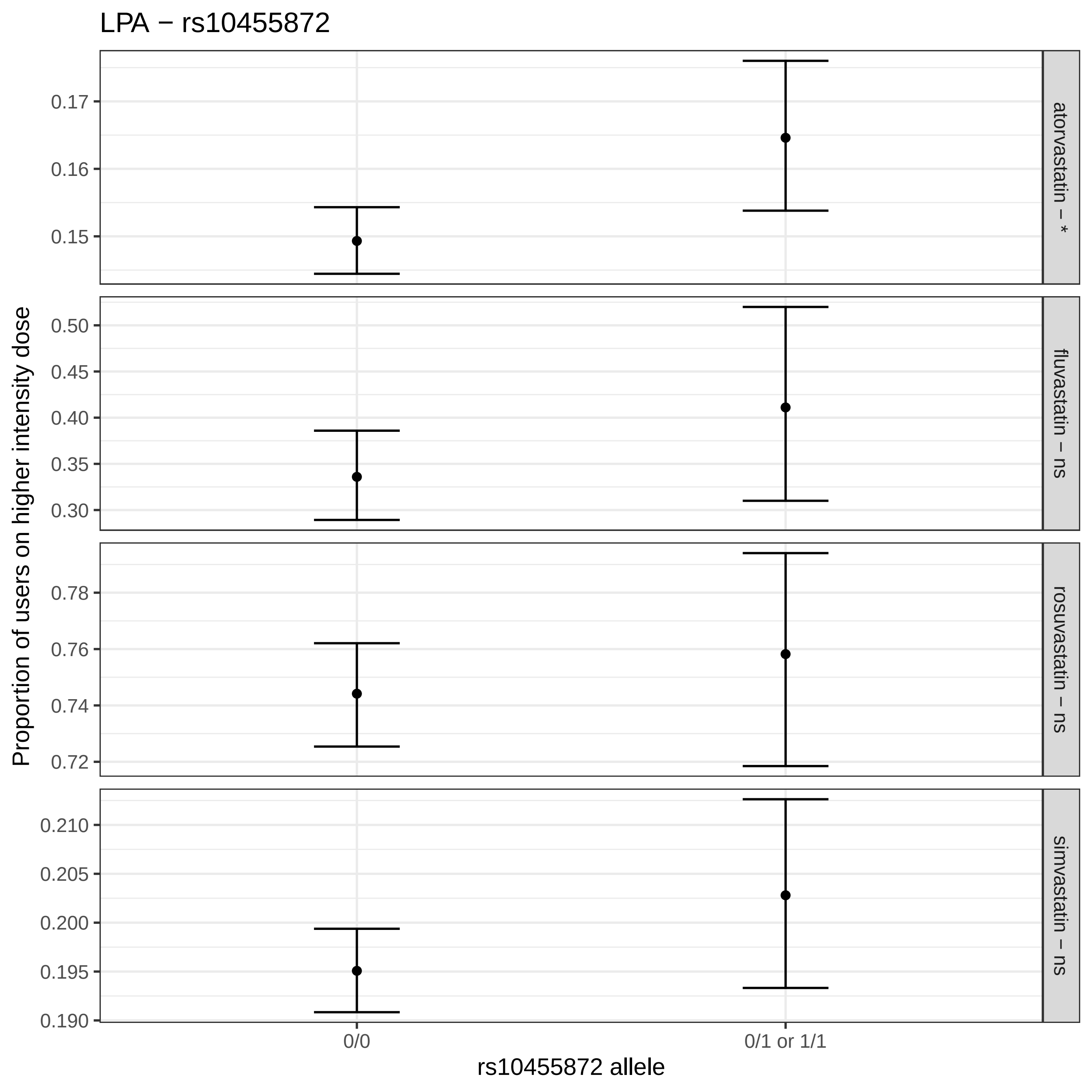

LPA − rs10455872
Proportion of users on higher intensity dose
0.17
0.16
0.15
0.50
0.45
0.40
0.35
0.30
0.78
0.76
0.74
0.72
0.210
0.205
0.200
0.195
0.190
atorvastatin − *
fluvastatin − ns
rosuvastatin − ns
simvastatin − ns
0/0
0/1 or 1/1
rs10455872 allele

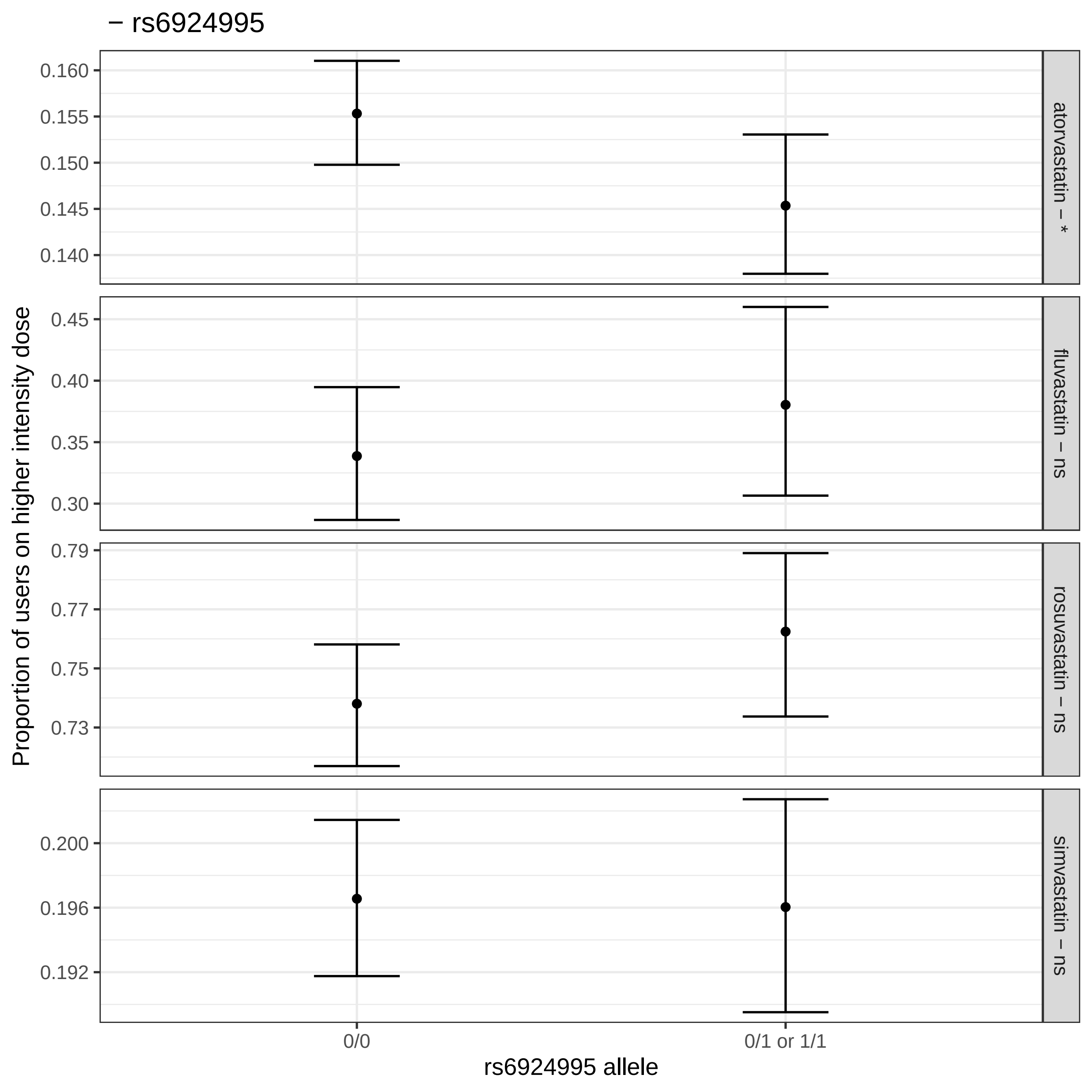
– rs6924995
Proportion of users on higher intensity dose
atorvastatin – *
fluvastatin – ns
rosuvastatin – ns
simvastatin – ns
0.160
0.155
0.150
0.145
0.140
0.45
0.40
0.35
0.30
0.79
0.77
0.75
0.73
0.200
0.196
0.192
0/0
0/1 or 1/1
rs6924995 allele

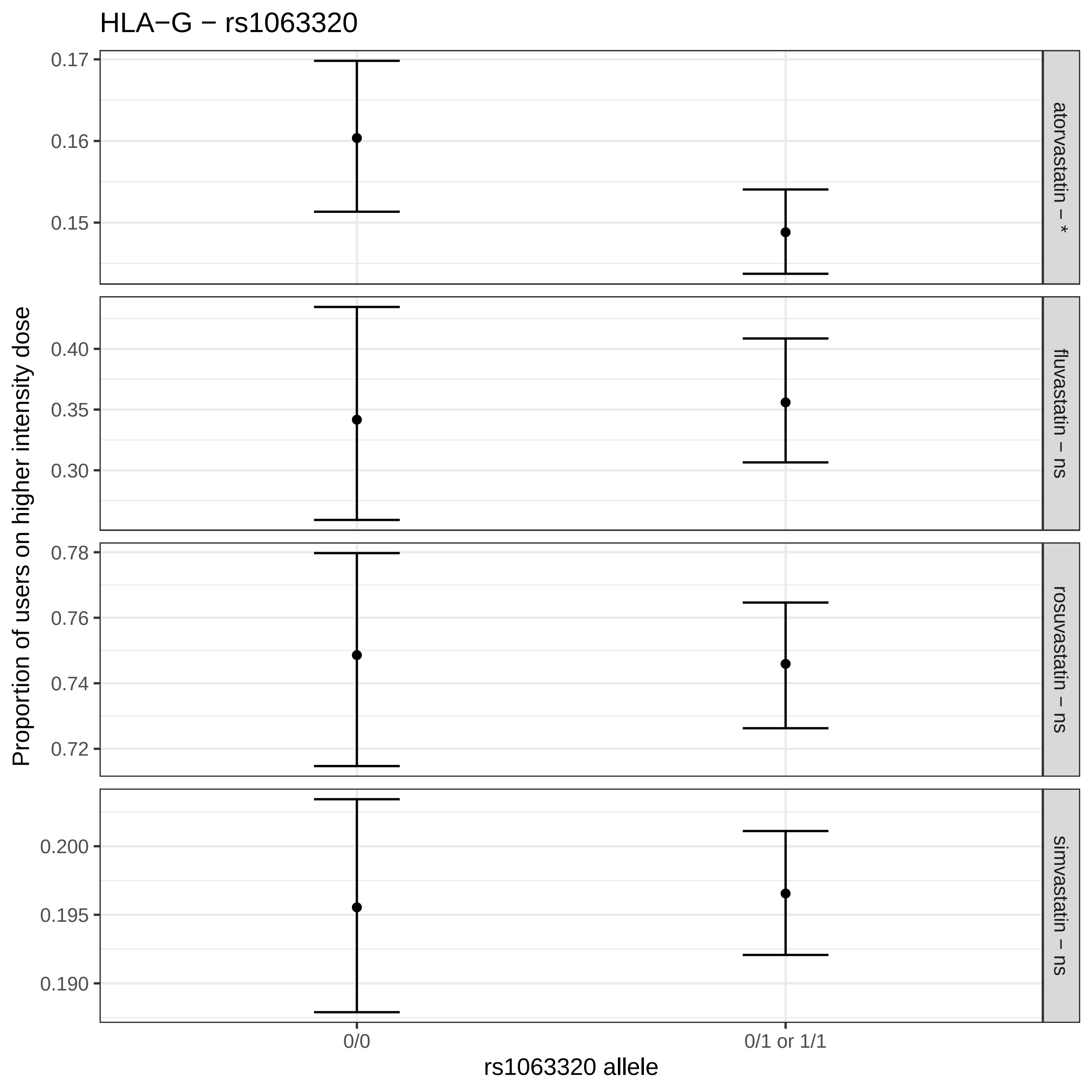
HLA−G − rs1063320
Proportion of users on higher intensity dose
atorvastatin − *
fluvastatin − ns
rosuvastatin − ns
simvastatin − ns
0.17
0.16
0.15
0.40
0.35
0.30
0.78
0.76
0.74
0.72
0.200
0.195
0.190
0/0
0/1 or 1/1
rs1063320 allele

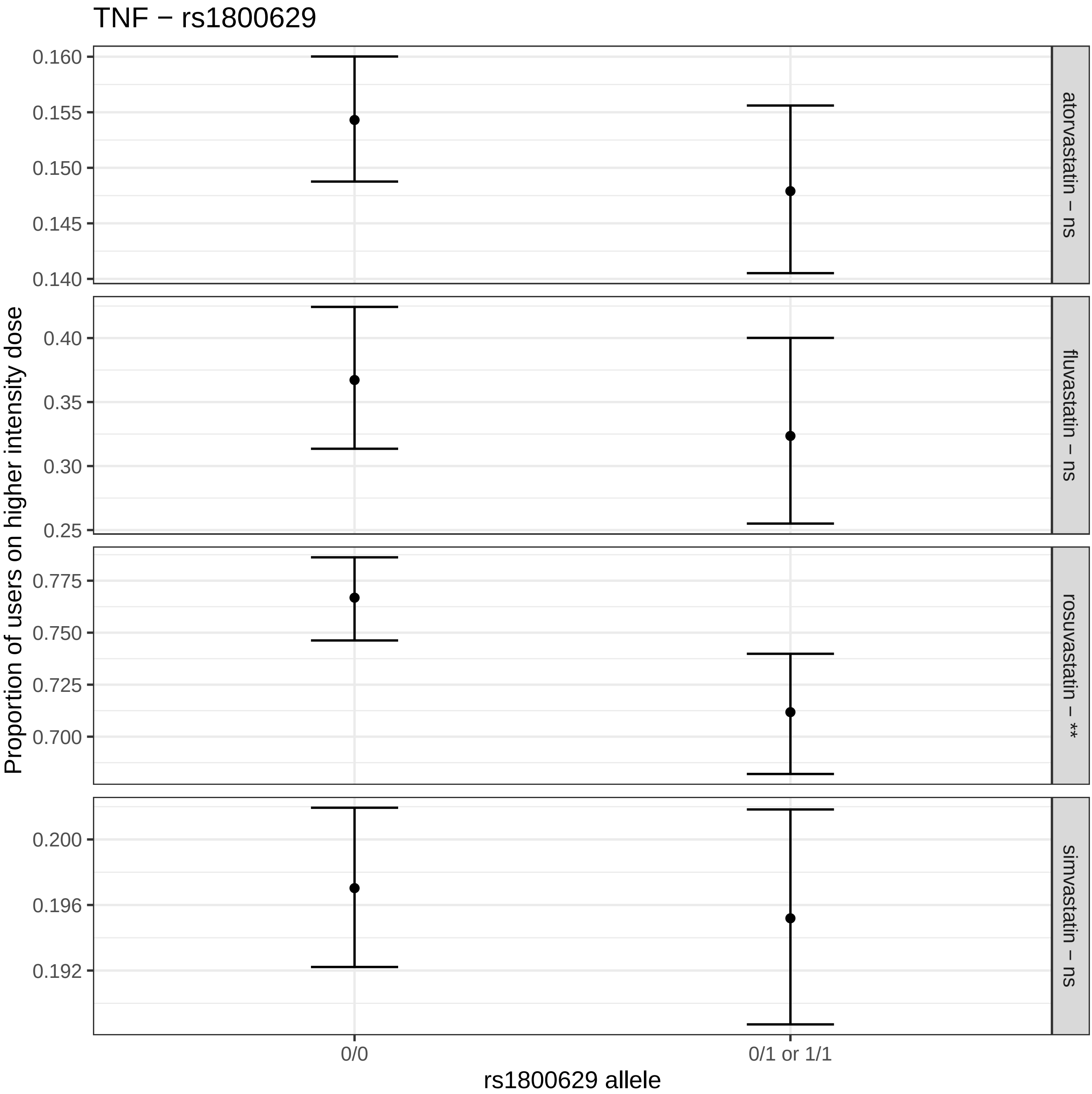
TNF − rs1800629
Proportion of users on higher intensity dose
0.160
0.155
0.150
0.145
0.140
atorvastatin − ns
0.40
0.35
0.30
0.25
fluvastatin − ns
0.775
0.750
0.725
0.700
rosuvastatin − **
0.200
0.196
0.192
simvastatin − ns
0/0
0/1 or 1/1
rs1800629 allele

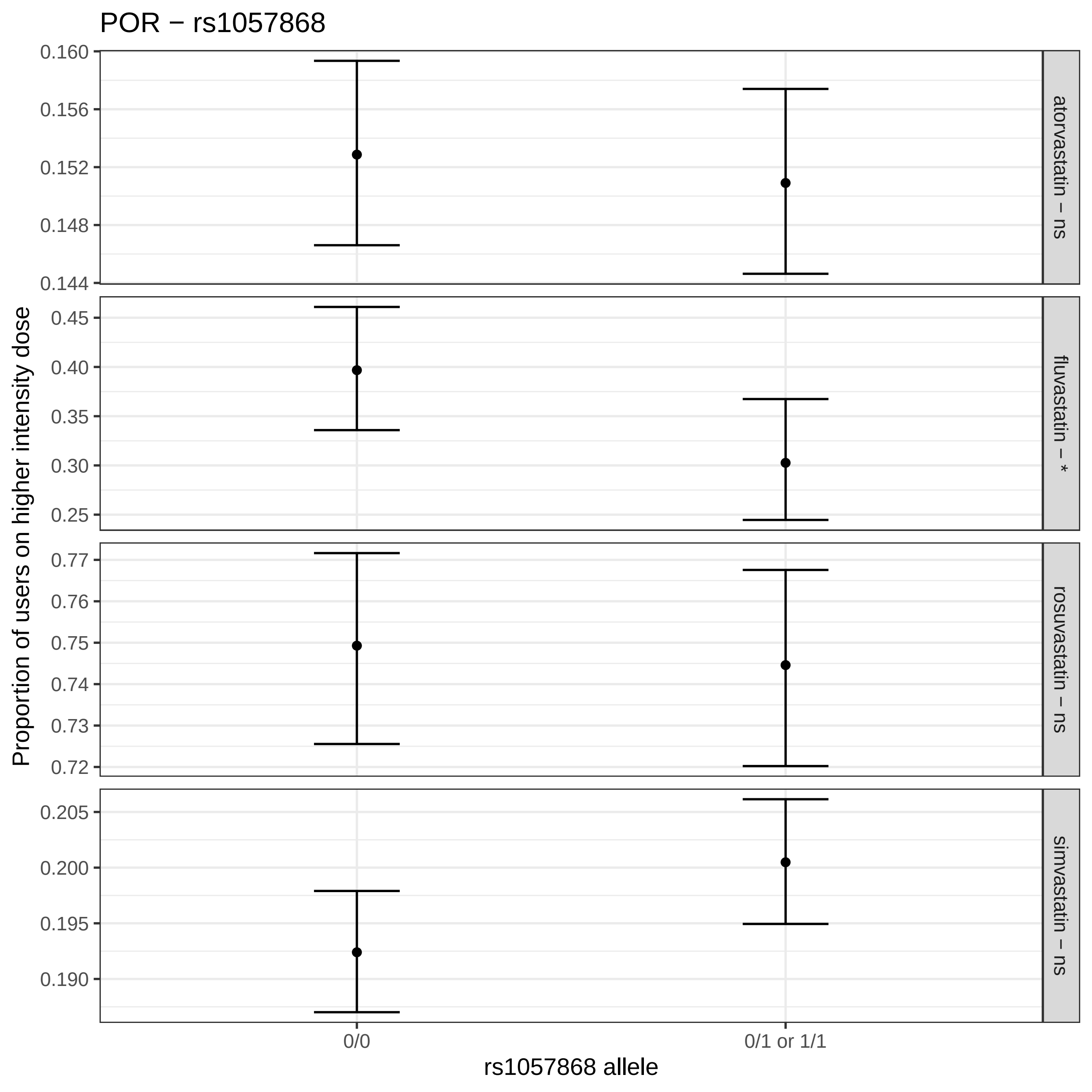

POR − rs1057868
Proportion of users on higher intensity dose
0.160
0.156
0.152
0.148
0.144
atorvastatin − ns
0.45
0.40
0.35
0.30
0.25
fluvastatin − *
0.77
0.76
0.75
0.74
0.73
0.72
rosuvastatin − ns
0.205
0.200
0.195
0.190
simvastatin − ns
0/0
0/1 or 1/1
rs1057868 allele

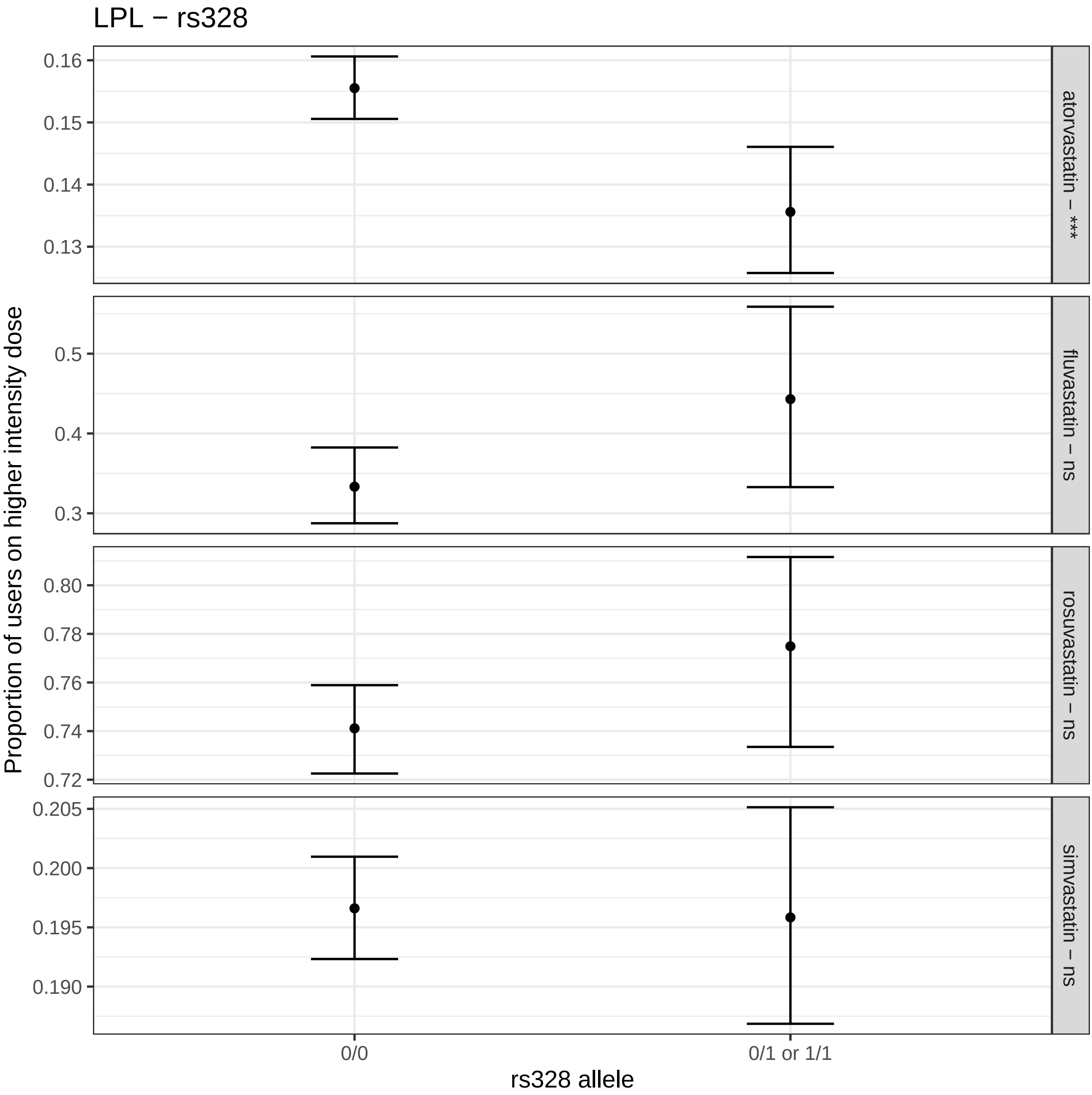

LPL − rs328
Proportion of users on higher intensity dose
0.16
0.15
0.14
0.13
atorvastatin − ***
0.5
0.4
0.3
fluvastatin − ns
0.80
0.78
0.76
0.74
0.72
rosuvastatin − ns
0.205
0.200
0.195
0.190
simvastatin − ns
0/0
0/1 or 1/1
rs328 allele

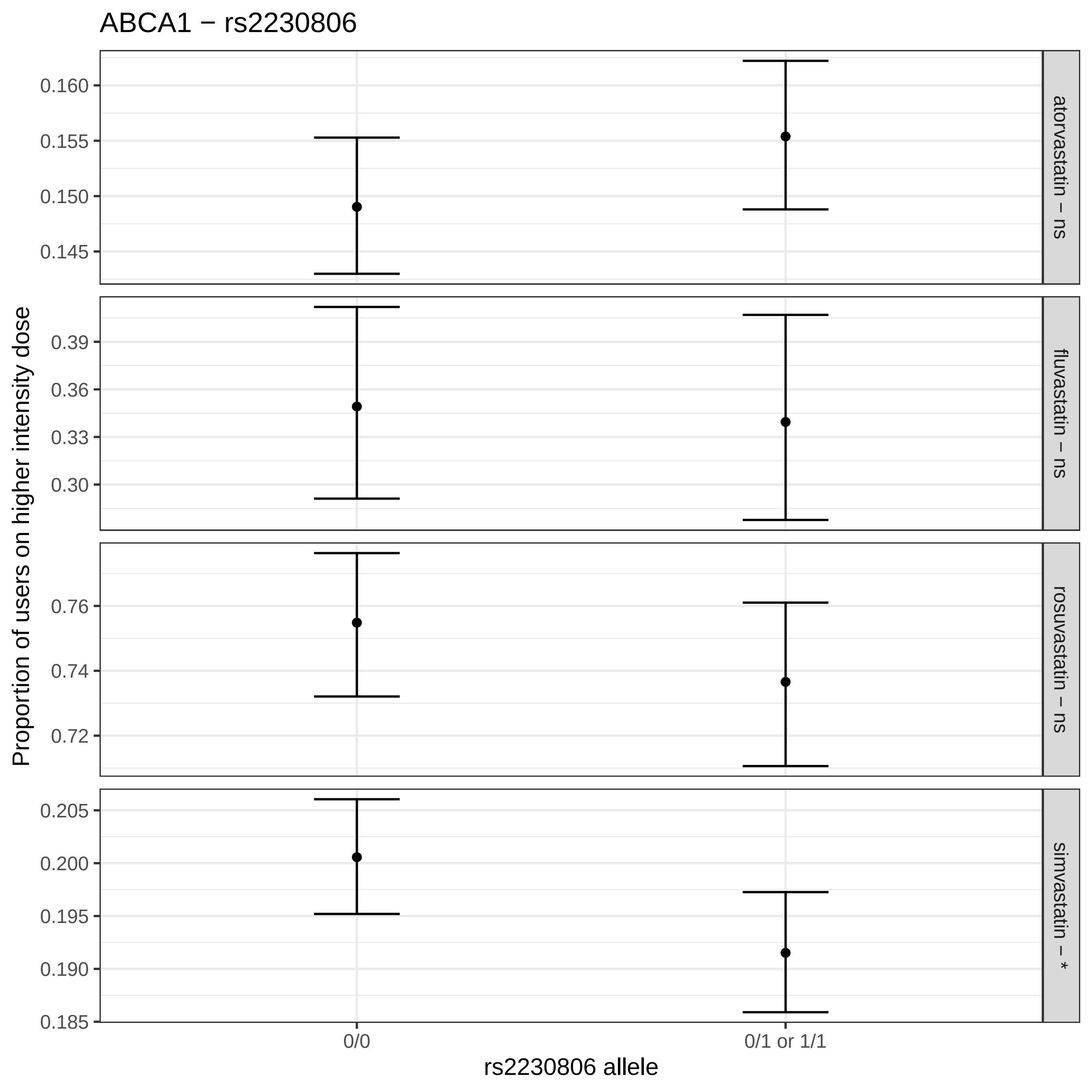

ABCA1 − rs2230806
Proportion of users on higher intensity dose
0.160
0.155
0.150
0.145
atorvastatin − ns
0.39
0.36
0.33
0.30
fluvastatin − ns
0.76
0.74
0.72
rosuvastatin − ns
0.205
0.200
0.195
0.190
0.185
simvastatin − *
0/0
0/1 or 1/1
rs2230806 allele
