## Supplemental Figures for "*LPA* and *APOE* are associated with statin selection in the UK Biobank"

**Supplementary Table 1.** Statins contained within the UK Biobank self-reported medication data and their associated medication codes. Each medication code, from Data-coding 4, represents an individual drug that was mapped to the generic name for the statin.

| Drug | Mapped UK Biobank Codes (Data-coding 4) |
|---|---|
| simvastatin | 1140861958,1140910652,1140881748,1141200040,1141188146,1140910654 |
| pravastatin | 1140888648,1140910632,1140861970 |
| fluvastatin | 1140888594,1140864592 |
| atorvastatin | 1141146234,1141146138 |
| rosuvastatin | 1141192410,1141187780,1141192414 |
| fenofibrate | 1140861954 |

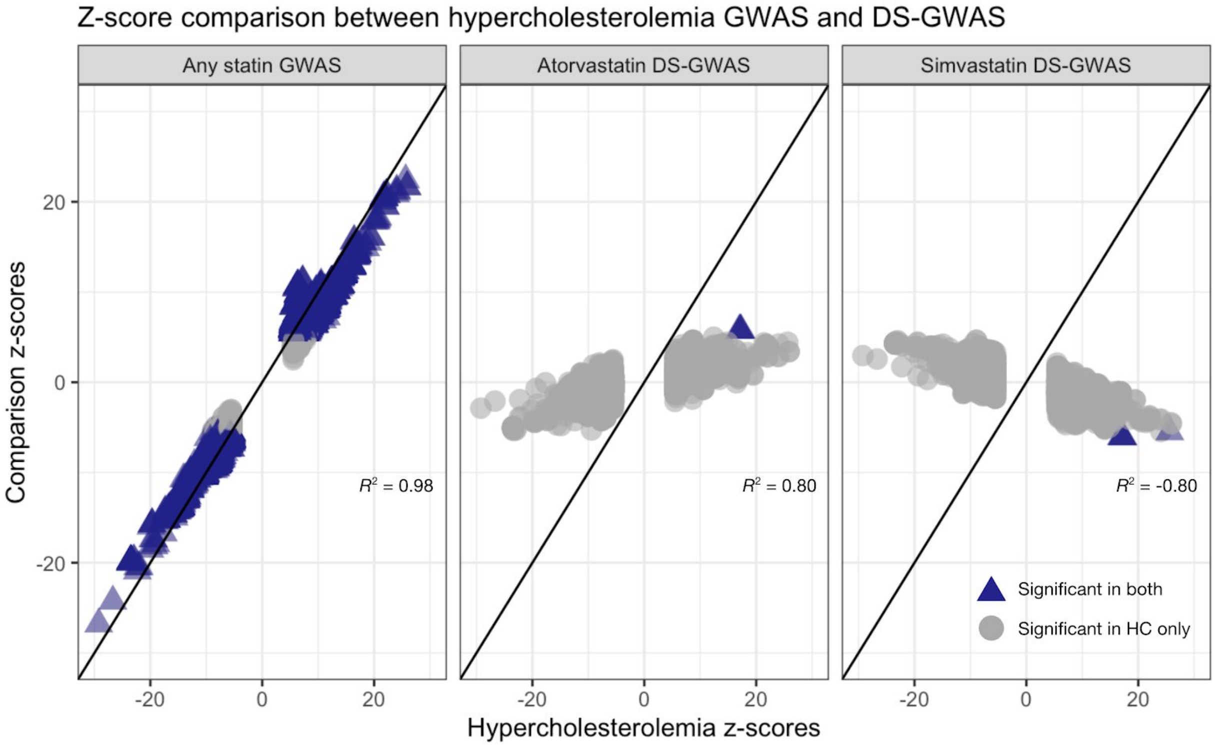


**Supplemental Figure 1.** Z-score correlations for high cholesterol GWAS and statin user GWAS (left) and high cholesterol GWAS and simvastatin DS-GWAS (right). Each figure shows all variants that were found to be significant in the high cholesterol GWAS, with the x-axis representing each variant's z-score in the high cholesterol GWAS and the y-axis representing the z-score in either the simvastatin user GWAS with controls being all subjects not on simvastatin (left) or the z-score from the simvastatin DS-GWAS (right). Blue triangles represent variants that were found to be significant in both analyses (corresponding to each individual plot), gray circles were only significant in the high cholesterol GWAS.
